# Interactions between ploidy and resource availability shape clonal interference at initiation and recurrence of glioblastoma

**DOI:** 10.1101/2023.10.17.562670

**Authors:** Zuzanna Nowicka, Frederika Rentzeperis, Richard Beck, Vural Tagal, Ana Forero Pinto, Elisa Scanu, Thomas Veith, Jackson Cole, Didem Ilter, William Dominguez Viqueira, Jamie K. Teer, Konstantin Maksin, Stefano Pasetto, Mahmoud A. Abdalah, Giada Fiandaca, Sandhya Prabhakaran, Andrew Schultz, Maureiq Ojwang, Jill S. Barnholtz-Sloan, Joaquim M. Farinhas, Ana P. Gomes, Parag Katira, Noemi Andor

**Affiliations:** Department of Biostatistics and Translational Medicine, Medical University of Łódź, Łódź, Poland; Icahn School of Medicine at Mount Sinai, New York City, NY, USA; Department of Integrated Mathematical Oncology, Moffitt Cancer Center, Tampa, FL, USA; Queen Mary University of London, London, United Kingdom; Cancer Biology PhD Program, University of South Florida, Tampa, FL, USA; Department of Molecular Oncology, Moffitt Cancer Center, Tampa, FL, USA; Department of Cancer Imaging and Metabolism, Moffitt Cancer Center, Tampa, FL, USA; Biostatistics and Bioinformatics, Moffitt Cancer Center, Tampa, FL, USA; Poznań University of Medical Sciences, Poznań, Poland; Quantitative Imagine Core, Moffitt Cancer Center, Tampa, FL, USA; Department of Cellular, Computational and Integrative Biology, University of Trento, Tento, Italy; Center for Biomedical Informatics & Information Technology and Division of Cancer Epidemiology and Genetics, National Cancer Institute, Bethesda, MD, USA; Department of Radiology, Moffitt Cancer Center, Tampa, FL, USA; Department of Mechanical Engineering, San Diego State University, San Diego, CA, USA

## Abstract

Glioblastoma (GBM) is the most aggressive form of primary brain tumor. Complete surgical resection of GBM is almost impossible due to the infiltrative nature of the cancer. While no evidence for recent selection events have been found after diagnosis, the selective forces that govern gliomagenesis are strong, shaping the tumor’s cell composition during the initial progression to malignancy with late consequences for invasiveness and therapy response. We present a mathematical model that simulates the growth and invasion of a glioma, given its ploidy level and the nature of its brain tissue micro-environment (TME), and use it to make inferences about GBM initiation and response to standard-of-care treatment. We approximate the spatial distribution of resource access in the TME through integration of in-silico modelling, multi-omics data and image analysis of primary and recurrent GBM. In the pre-malignant setting, our in-silico results suggest that low ploidy cancer cells are more resistant to starvation-induced cell death. In the malignant setting, between first and second surgery, simulated tumors with different ploidy compositions progressed at different rates. Whether higher ploidy predicted fast recurrence, however, depended on the TME. Historical data supports this dependence on TME resources, as shown by a significant correlation between the median glucose uptake rates in human tissues and the median ploidy of cancer types that arise in the respective tissues (Spearman r = -0.70; P = 0.026). Taken together our findings suggest that availability of metabolic substrates in the TME drives different cell fate decisions for cancer cells with different ploidy and shapes GBM disease initiation and relapse characteristics.

## Introduction

The majority of human cells are diploid. A whole-genome duplication (WGD) transforms a diploid cell into a tetraploid state. Tetraploid cells are genomically unstable and the population goes on to become aneuploid – a hallmark of all cancers, including glioblastoma (GBM). In GBM, ∼14% of tumors show evidence of at least one WGD event in their evolutionary history^1^. WGD tumors typically evolve to a ploidy of approximately three by the time of tumor detection, in contrast to non-WGD tumors, which tend to remain near-diploid. Cancers differ substantially in their ploidy levels^1–3^. Incidence of tumors with evidence of past WGD events ranges from less than 10% in Non-Hodgkin’s Lymphoma to almost 60% in germ cell cancer^1^. If a WGD subclone expands, it consistently does so early in tumor evolution^1,4^, when cell density is still low and competition for resources is comparatively weak – an observation confirmed for several tumor types^12,16,17^. This suggests that in tumors with no evidence of WGD, pre-malignant WGD cells have likely been driven to extinction by co-existing non-WGD cells. Indeed, longitudinal studies of GBM as well as analyses of spatially mapped tumor biopsies found no evidence for significant selection pressure during progression, even in response to treatment, suggesting selection in GBM happens early, before tumor detection. The outcome of the competition between pre-malignant WGD and non-WGD cells has an important influence on the ploidy of the tumor at the time of detection. Ploidy in turn predicts sensitivity to multiple classes of therapies in-vitro, including cytotoxic agents and targeted therapy^5^. Considering that WGD+ cells need more resources, especially in the form of phosphate, glucose and oxygen to replicate an increased amount of DNA, and synthesize more RNA and proteins^6^, it is likely that resource availability in the brain tissue micro-environment (TME) shapes the outcome of the competition between a WGD cell and its diploid ancestor^7,8^. This TME context early during gliomagenesis may thus have a profound impact on the tumor’s clonal composition, invasiveness and therapy response. Cancer cell ploidy, through its significance for resource requirements on the one hand and for evolvability and plasticity on the other, could potentially play a key role in therapy response.

While advances in next generation sequencing technologies have facilitated precise characterization of a tumor’s clonal composition, including its WGD status, characterization of the TME has been mostly focused on its immune cell composition. Spatio-temporal TME variability in terms of cell-extrinsic resources has not been well described, hampering the study of its’ role in GBM initiation and recurrence. Based on the limited evidence available, it is now clear that physiologic oxygen concentrations vary across different brain regions, and intracerebral tumor location is a strong predictor of the degree of hypoxia in the tumor^9^. Cerebral oxygenation is compromised in older humans and hypoxia plays a central role in brain aging. Aging brains also have higher incidence of polyploidy (Pearson r=0.80; P=0.031)^10^, and glucose levels^11^ as well as decreased brain tissue stiffness^12^. GBM prognosis is considerably worse among the elderly, with patient age being a strong and independent predictor of worse survival^13^. Together these studies suggest that the intracerebral point of origin of the first pre-malignant GBM cells is an important candidate predictor of the route of tumor progression, and emphasize the value of quantifying spatio-temporal variation in resource access.

Our manuscript is structured into three parts. In the first part, under methods, we present a mathematical model (Stochastic State-Space Model of the Brain – S3MB) that simulates the growth and invasion of a glioma, given its ploidy and the composition of the TME where it originates. In the second part, we derive three atlases of the brain TME by quantifying its resource availability with an integrated in-silico, multi-omic and imaging analysis of healthy volunteers and primary and recurrent GBM. Finally, in the third part, covered by the last two result sections, we use S3MB in the context of the TME information and ploidy composition to make inferences about GBM initiation and recurrence. Hereby we address the question whether resource access during early pre-malignancy can shape ploidy composition, late-stage tumor progression and therapy response.

## Methods

### S3MB: a scalable mathematical model of glioma growth

Several computational models that simulate GBM growth as a function of the brain microenvironment, using reaction diffusion PDEs, have been previously developed^14,15^. We build upon this work to develop a new model whose assumptions in part overlap with those made by the previously developed proliferation-invasion-hypoxia–necrosis– angiogenesis (PIHNA) model of glioma growth^16^. The PIHNA model partitions the tumour cells into subpopulations based on histological characteristics such as normoxia, hypoxia, necrosis and angiogenesis, modeling interactions between them. What prior mathematical models of GBM invasion have been grappling with, however, are three challenges: the integration of multiple data types, the impact of intra-tumor heterogeneity and evaluation of the predictive performance of models in patient-derived data^15^. To address these three challenges, we developed a Stochastic State-Space Model of the Brain (S3MB). Our model discretizes the brain into voxels, and the state of each voxel is defined by brain tissue stiffness, oxygen, glucose, vasculature, dead cells, migrating cells and proliferating cells of various ploidies, and treatment conditions such as radiotherapy and chemotherapy with temozolomide (TMZ). Well established Fokker-Planck partial differential equations govern the distribution of resources (oxygen, glucose) and cells across voxels^17^. However, we modified these equations as described below to simplify the model and rapidly track the temporal changes in live cells of ploidy *i* (*n*_*i*_), dead cells (*N*_*d*_), oxygen (*N*_*o*_) and glucose (*N*_*g*_) as well as states of vascular supply (*V*) and brain tissue stiffness (*S*):

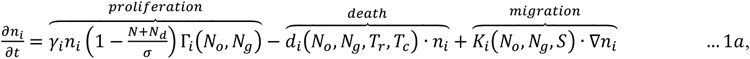

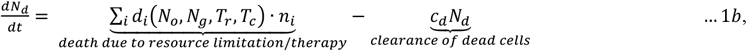

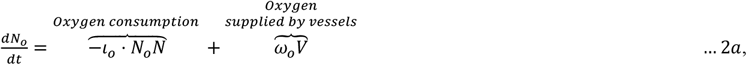

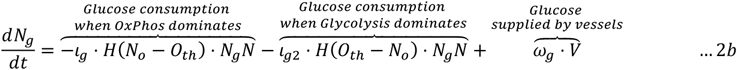

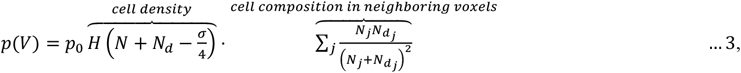

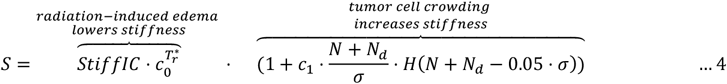

Interacting elements of the S3MB model are described in Figure 1 and a detailed description of all model components, simplifcations and assumptions is given in Supplementary section 1. Cell migration (*K*) is determined by brain tissue stiffness and by glucose and oxygen availability. In addition to influencing cell migration, oxygen availability has several downstream effects on the behaviour of the system: it impacts both chemo- and radiation therapy induced cell death (via parameters α_*Tchemo*_ and *r*^*\**^ respectively) as well as switching between oxidative phosphorylation and glycolysis (further referred to as metabolic state). The metabolic state in turn also determines the rate at which cells proliferate (Michaelis menten constants α_g_ or β_g_), die (Michaelis menten constants α_*d*_ or α_*d*_) and consume glucose (rates l_g_ and l_g2_). Both dead and live cell concentrations in focal and neighboring voxels determine the probability of vascular recruitment (*p*(*V*)). Stiffness of the brain tissue within each voxel (*S*) in conjunction with available resources dictates the migration potential of cells^18^ and is influenced by both, treatment and tumor cell density. S3MB was implemented in MATLAB. At each specified time, cell and resource densities are updated, whereby cell death happens first, followed by proliferation and migration. While cell migration, growth and death as well as overall resource utilization is obtained deterministically, vasculature recruitment to specific voxels is stochastic. The outcomes of the model strongly depend on the initial location of tumor cells, the initial tumor composition in terms of cell ploidy, resource perfusion within the brain tissue as well as brain tissue stiffness. We can simulate tumor growth over 365 days within 10 seconds, as well as include specific treatment regimens and other clinically relevant information within our model simulations.

**Figure 1:**
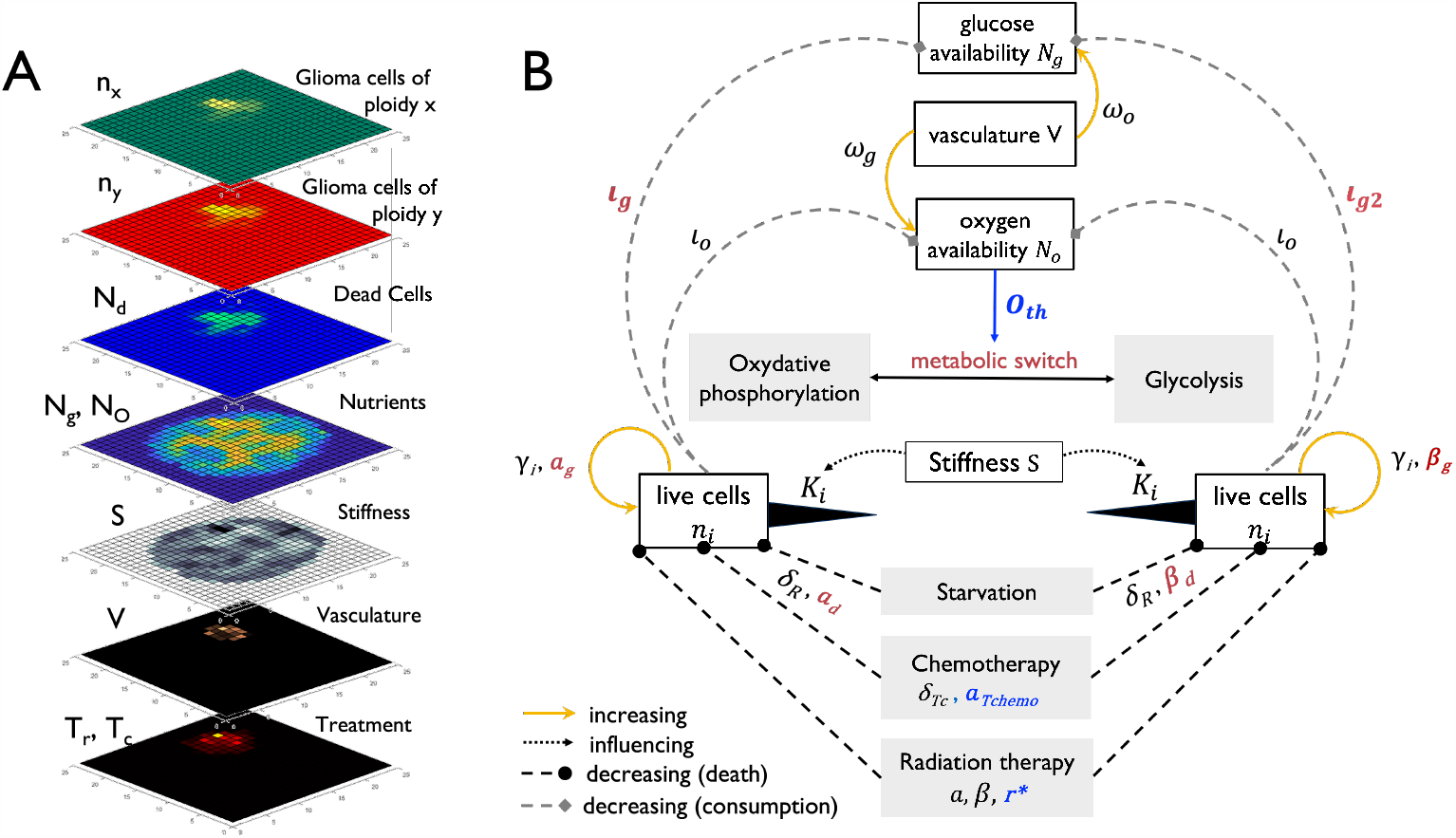
Stochastic State-Space Model of the Brain (S3MB) – (**A**) Model Elements. S3MB simulates glioblastoma (GBM) growth within the brain tissue while tracking oxygen and glucose as nutrients, nutrient-dependent treatment efficiency, stiffness and glucose dependent cell migration, dynamically altering clonal composition and stochastic recruitment of vasculature. (**B**) Interactions between model compartments (white boxes) in the context of different metabolic and/or therapeutic environments (gray boxes). Parameters depending on either the metabolic switch or on oxygen availability are marked in red and blue respectively.

### In-vitro quantification of cell migration in response to glucose deprivation

The selective forces that govern the outcome of the competition between a WGD+ cell and its diploid ancestor set a crucial branching point in the tumor’s evolutionary history. Depending on this outcome, the cellular phenotypes that follow will differ. For proliferating cells, migration is a first response to suboptimal resource levels^38–40^. Prior studies suggest that a proliferating GBM cell will start migrating if its environment provides insufficient resources to continue proliferation and that it will cease migrating either when it starts dying or when it can continue proliferation^41^. I.e.: Cells migrate to sustain their proliferation. Because high ploidy cells need more resources than low ploidy cells in order to proliferate, we hypothesized that a WGD+ cell growing next to a WGD-cell in the same resource scarce environment, will seize proliferating sooner and start migrating sooner, as it is further away from its energetic optimum. In support of this hypothesis, recent work has shown that WGD+ cells have increased metastatic potential compared to their near-diploid ancestors^42^ and gain a mesenchymal phenotype^43^. To test this hypothesis, we used a two-chamber culture system (Clearview Cell Migration plate, see also Supplementary section 3.2). Whereas glucose and FBS concentrations were kept constant in the bottom chamber at 25 mM, we varied glucose and FBS concentrations in the upper chamber from 0.25-25 mM and 0.1-10% respectively. Isogenic near-2N and near-4N cells (see Supplementary section 3) were seeded in the upper chamber and allowed to migrate into the lower chamber through 8 um pores. The plate was imaged using the chemotaxis assay in a Sartorius Incucyte and images were taken every 2h for 48h (Supplementary Figure 7A). We processed 6000 raw images from the Incucyte transwell assay (80 wells; 25 times points per well; and three channels Phase, Top, and Bottom). To create a training and test dataset, we cropped two random regions of 200 × 200px from each frame selected. We used the image classification and segmentation software *ilastik*^44^ to classify cells as “top alive”, “bottom alive”, and “top dead” (Supplementary Figure 7B; see also Supplementary section 3.3). Comparison of the pipeline’s output on the test dataset with manual counts confirmed a good agreement in the number of detected cells per each of the three classes (Supplementary Figure 8).

We developed an ODE model of cell migration in response to glucose deprivation (Supplementary Figure 7C and Supplementary section 3.4), which describes the temporal evolution of these three cell compartments: Top alive (*T*_*a*_), bottom alive (*B*_*a*_), and top dead, (*T*_*d*_). In this model, *T*_*a*_ cells can proliferate at a rate *λ*_*d*_, migrate to the bottom chamber at a rate μ and become *B*_*a*_ cells, or die at a rate δ and become *T*_*d*_ cells (see also Supplementary section 3.4). To infer how glucose concentration in the top chamber affects migration, proliferation, and death rate of *T*_*a*_ cells, we fitted the ODE model to the data from the five top chamber glucose concentrations (0.1, 0.5, 1, 5 and 25 mM; Supplementary Figure 9). While proliferation rate was minimal in all glucose concentrations (presumably because of the limited FBS concentration in the top chamber), death-rate increased with decreasing glucose concentration (4N: Pearson r = -0.94; p-value<<1E-6; 2N: Pearson r = -0.74; p-value=1.8E-4). A similar non-monotonic relation between glucose concentration and migration rate was observed for both 2N and 4N cells, with 4N cells migrating faster than 2N cell under all glucose concentrations (Supplementary Figure 7D). We conclude that the bimodal cell migration formulated in eq. *s6* of S3MB accurately reflects cell behaviour and that these in-vitro results can be used to infer its parameters (*a*_*mig*_ and *b*_*mig*_ ; Supplementary Figure 7D).

### Whole-slide digital pathology for estimation of vascular throughput and carrying capacity in GBM

We used three H&E stained samples obtained from the primary and recurrent GBM of the same patient to approximate glucose generation rate (additional glucose supplied by recruited vasculature). H&E digital slides were analyzed and annotated by a histopathologist (Supplementary section 4.2). After identifying multiple tumor core (TC) and infiltrating zone (IZ) as circular regions of interest (ROI), we quantified tumor cells using QuPath software^45^ and manually marked vascular architecture of each ROI (**Figure 2**A,B). By calculating the average cell density over ROIs we approximated carrying capacity *S* for S3MB as σ = *max* (*Nx*), *x* ∈ *ROI*, with the result (σ ≈ 15,448 *cells/voxel*) approximately matching estimates from literature^46^ (∼20,000*cells/voxel*).

**Figure 2:**
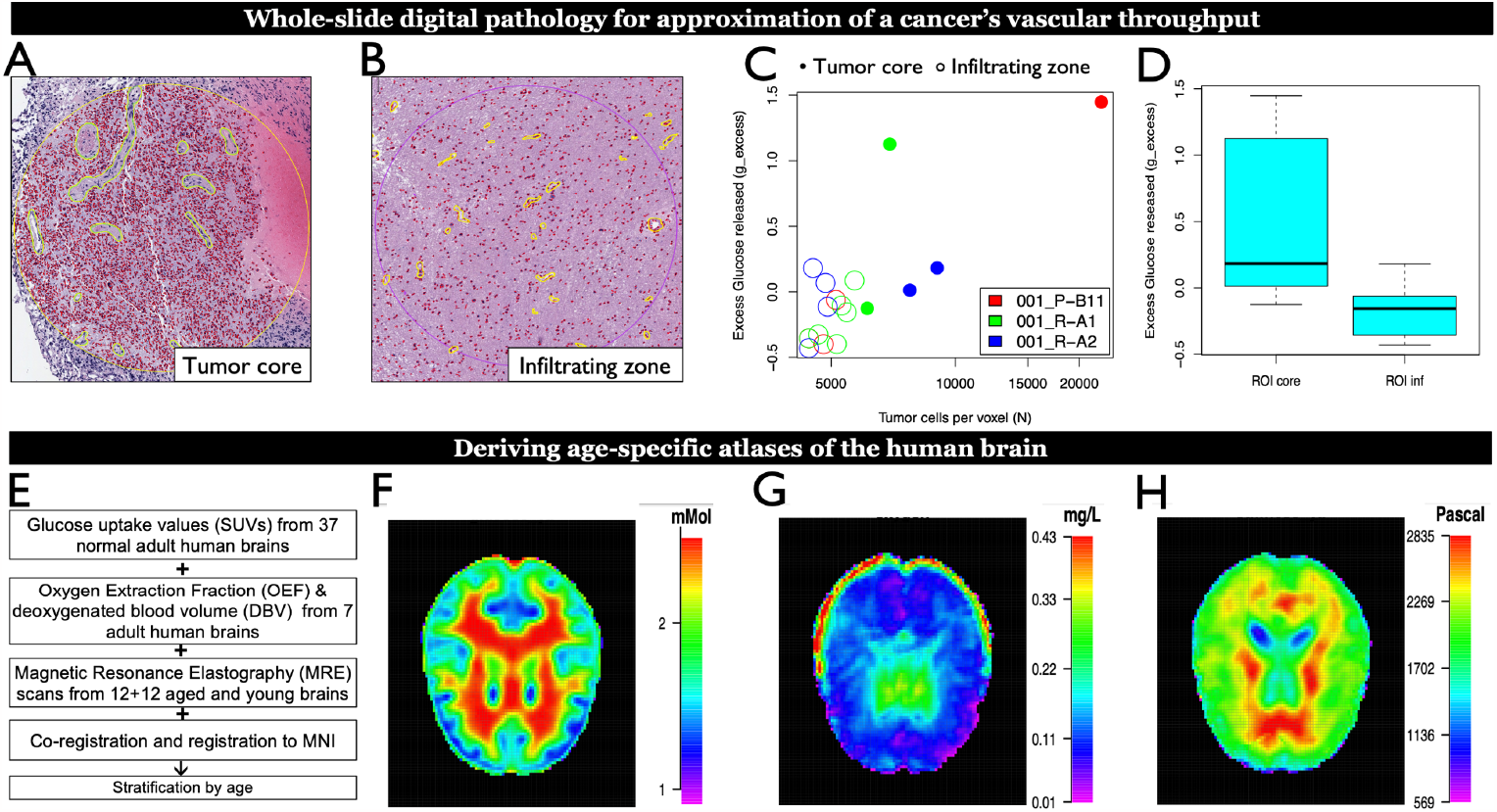
Multi-modal imaging analysis to characterize GBM and brain tissue microenvironments. (A-D) Whole-slide digital pathology analysis and results. Three whole-slide images obtained from the primary and recurrent surgery of a GBM patient were analyzed for representative tumor core regions (A) and tumor infiltrating zones (B).For each region, cells were segmented, and vasculature was manually labeled by a pathologist. (C) The total area occupied by vasculature was used to approximate the excess Glucose released (y-axis) and was found to correlate to tumor cell density (x-axis). (D) Tumor core regions are thus assumed to release more excess Glucose than tumor infiltrating zones. This is a direct consequence of the higher vascular density in the former. (E-H) Medical imaging analysis and results. (E) Overview of analyzed datasets, resulting in a glucose atlas (F), an Oxygen atlas of the human brain (G), and a brain tissue stiffness atlas (H).

Next, we calculated the total area occupied by vasculature (*V*_*a,i*_) for each ROI, *i*, and found that both cell and vascular density were higher in TC than in IZ regions (Wilcoxon rank-sum test: p-value = 0.019; **Figure 2**C,D). To estimate the glucose supply from recruited vasculature, we assume further that the glucose generation rate is directly proportional to vascular density and that, since vasculature is denser in regions adjacent to the tumor, the average vascular density across all IZ ROIs approximates that of normal brain regions. Following the above assumptions, the fold difference in vascular density between tumor and normal brain regions yields an approximation of additional glucose generated per each ROI:

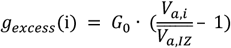

where *G*_*0*_ is the average glucose generated per day by physiologic cerebral vascular supply (Supplementary section 4.2). We approximated S3MB parameter ω_*g*_ as the median excess glucose produced by *TC* regions: 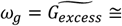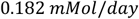. We repeated the approach analoguously to approximate median excess oxygen produced by *TC* regions: 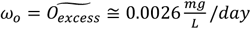 (Supplementary section 4.2).

### Medical imaging for derivation of glucose-, oxygen- and brain tissue stiffness atlases

We analyzed a total of 68 subjects from three public datasets to derive glucose uptake, oxygen and tissue stiffness atlases of the human brain. Briefly, for each dataset and subject, functional and anatomical scans were co-registered prior to registration of the anatomical scan to the Montreal Neurological Institute (MNI) standard space.

We derived an atlas of glucose levels from a public database of 37 healthy adult human brains. It is comprised of [^18^F]FDG PET, T1 MRI, FLAIR MRI, and CT images for each brain^47^. Following co-registration and registration to MNI space, PET images were normalized to obtain SUV and Standard Uptake Value ratio (SUVr) images at a ≤ 5% risk of Type 1 error^47^. These were used here and stratified by the patient’s age. To approximate glucose levels, we assumed SUVr values are inversely correlated to glucose uptake^8 48,49^ and that glucose concentration in the brain ranges between 1.0-2.7 mMol^50^. This was achieved through reflection of SUVr values, followed by scaling to the known range of brain glucose concentrations: *Ng* = *scale* (− *SURr*, [1.0, 2.7 *mMol*]). Since SUVr values are dimensionless, the scaling also assigns a unit to the resulting estimate *Ng*.

An atlas of oxygen level was calculated from quantitative BOLD (qBOLD) data from seven volunteers, downloaded from the Oxford Research Archive site^51^. All image processing was performed with Quantiphyse^52^. The mean reversible transverse relaxation rate (T2), the oxygen extraction fraction (oef) and the deoxygenated blood volume (dbv) were inferred with Bayesian modelling using the qBOLD widget in Quantiphyse^53^. The method deconvolutes three major confounding effects of qBOLD: (i) Cerebral Spinal Fluid (CSF) signal contamination; ii) Macroscopic magnetic field inhomogeneity and iii) Underlying transverse T2 signal decay^53^. Following registration, an atlas of dissolved oxygen concentration (dO2) was derived from oef, in units of mg/L as:

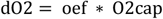

where O2cap = 0.4655 mg/L is the typical average oxygen content of fully oxygenated capillaries^54^. Tissue oxygenation is a complex process influenced by various factors – above equation provides a simplified representation of it. It approximates the oxygen content in brain tissue, under the assumption that the extracted oxygen is retained by the tissue.

For the stiffness atlas, magnetic Resonance Elastography (MRE) scans were obtained from a database of 12+12 aged and young brains generated at the University of Edinburgh, UK^12^. MRE scan was were co-registered to the T1 scan via rigid body transformation. The MRE scans were then transformed to MNI space by the same protocol as the PET scans.

#### Modeling the effect of surgery on oxygen, glucose and stiffness atlas

Because stiffness, glucose, and oxygen maps all share the same MNI coordinate system as the tumor cell density maps (see Supplementary section 4.3), we can model the effect of surgery on these environmental factors. Let *CAV* denote the resection cavity map after a given surgery and *S, N*_*g*_, *N*_*o*_ are the stiffness, glucose, and oxygen maps. We assume tumor resection decreases stiffness as well as perfusion in the surgery-affected brain region (due to lack of a fractal pattern in blood supply provided by vasculature), as follows:

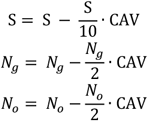

Detailed protocols describing how each of the three atlases was derived is available in Supplementary section 2. By registering a given GBM patient’s MRI onto the same MNI coordinate system, the three atlases allow for approximation of the microenvironment in the vicinity of the tumor.

## Results

### Characterization of resource distributions in the glioma and healthy brain tissue environment

S3MB is highly efficient in simulating GBM growth within the brain tissue, combining several unique features including stochastic recruitment of vasculature, independent tracking of oxygen and glucose as nutrients, nutrient dependent treatment efficiency, stiffness dependent cell migration, and dynamically altering clonal composition. An overview of all 25 model parameters is given in Table 1. Parameters representing resource-driven phenotypic transitions and resource concentrations play an important role in our model, yet have to our knowledge not been well described. We performed a multi-modal data analysis for quantification of resources in the brain tissue environment.

**Table 1:** Overview of model parameters. Parameters were inferred from literature, in-vitro results and from patient data. For fitted parameters, shown here are only initial (literature-informed) values . The source types are as follows: Level 1 – literature or experimental data specific to glioblastoma (GBM), Level 2 – literature or experimental data from other cancer type or cell line. A voxel is defined as a square of 4 mm^2^.

|  | Name | Value | Value range | Unit | Description | Source | Source type |
| --- | --- | --- | --- | --- | --- | --- | --- |
| 1 | $\gamma$ | 0.063 | 0.063-0.303 | cells/day | proliferation rate | Rew et al., 2000<sup>19</sup> | Level 1 |
| 2 | $\alpha_d$ | 0.0025 | 0.0007-0.034 | mg/L | oxygen at which cell survival is half-max. | Papandreou et al., 2005<sup>20</sup> | Level 2 |
| 3 | $\beta_d$ | 1.00 | 0-2.9 | mM | glucose at which cell survival is half-max. | Maeda et al., 2020<sup>21</sup> | Level 2 |
| 4 | $\alpha_g$ | 0.003 | 0.0007-0.034 | mg/L | oxygen at which proliferation is half-max. | Papandreou et al., 2005<sup>20</sup> | Level 2 |
| 5 | $\beta_g$ | 2.949 | 0.1988-5.7 | mM | glucose at which proliferation is half-max. | López-Meza et al., 2016<sup>22</sup> | Level 2 |
| 6 | $a_{mig}$ | 0.53/0.34 | N/A | 1/mM | biphasic migration rate constant of 2N/4N cells | in-vitro: Supplementary section 3 | Level 2 |
| 7 | $b_{mig}$ | 0.57/0.95 | N/A | 1/mM | biphasic migration rate constant of 2N/4N cells | in-vitro: Supplementary section 3 | Level 2 |
| 8 | $v_{max}$ | 0.36 | 0.17-0.50 | voxels/day | top migration speed based on cell type | Diao et al., 2019<sup>23</sup> | Level 1 |
| 9 | $\sigma$ | 15,448 | N/A | cells/voxel | carrying capacity | H&E: Supplementary section 4.2 | Level 1 |
| 10 | $Stiff_0$ | 2930 | 2530-2930 | pascals (Pa) | max tissue stiffness | Weickenmeier et al., 2016<sup>24</sup> | Level 1 |
| 11 | $c_0$ | 0.987 | N/A | dimensionless | radiation-induced change of stiffness | Kuroiwa et al., 1997<sup>25</sup>; Wang et al., 2021<sup>26</sup>; Supplementary section 1 | Level 1 |
| 12 | $c_1$ | 6.6792 | N/A | dimensionless | change in stiffness from remodelling by tumor cells | Isomursu, et al., 2022<sup>27</sup> | Level 2 |
| 13 | $c_2$ | -1.38E-8 | N/A | 1/Pa<sup>2</sup> | quadratic parameter for optimal tissue stiffness for directed cell migration | Isomursu et al., 2022<sup>27</sup> | Level 2 |
| 14 | $c_3$ | 2.35E-3 | N/A | 1/Pa | linear parameter for optimal tissue stiffness for directed cell migration | Isomursu et al., 2022<sup>27</sup> | Level 2 |
| 15 | $c_d$ | 0.63 | 0.48-0.78 | frac. cells/day | clearance rate of dead cells | Klöditz et al., 2019<sup>28</sup>; Khan et al., 2023<sup>29</sup> 17/10/2023 06:21:00 | Level 2 |
| 16 | $\delta_R$ | 0.171 | 0.053-0.209 | frac. cells/day | starvation shrinkage rate | Stensjøen et al., 2015<sup>30</sup> and Rew et al.<sup>19</sup> | Level 1 |
| 17 | $p_0$ | 0.2 | 0-0.40 | dimensionless | max angiogenesis probability | Chaplain et al., 2006<sup>31</sup> | Level 2 |
| 18 | $\omega_g$ | 0.18204 | N/A | mM/day | add. Gluc. generated by new vasculature | H&E: Supplementary section 4.2 | Level 1 |
| 19 | $\omega_o$ | 0.00257 | N/A | mg/L/day | add. Oxygen generated by new vasculature | H&E: Supplementary section 4.2 | Level 1 |
| 20 | $\alpha$ | 0.102 | -0.05-0.25 | 1/Gy | alpha in the radiation therapy induced cell death equation | Barazzuol et al., 2010<sup>32</sup> | Level 1 |
| 21 | $\beta$ | 0.008 | -0.012-0.026 | 1/Gy<sup>2</sup> | beta in the radiation therapy induced cell death equation | Barazzuol et al., 2010<sup>32</sup> | Level 1 |
| 22 | $r^*$ | 3.00 | 2.1-3.7 | dimensionless | oxygen enhancement ratio | Murray et al., 2003<sup>33</sup> | Level 1 |
| 23 | $a_{Tche mo}$ | 1.15 | N/A | mg/L | oxygen at which TMZ-shrinkage rate is half-max. | Musah-Eroje et al., 2019<sup>34</sup> | Level 1 |
| 24 | $\delta_{Tc}$ | 0.0196 | N/A | frac. cells/day | TMZ-induced shrinkage rate (max) | Stupp et al., 2005<sup>35</sup>; Elazab et al., 2017<sup>36</sup>; Barazzuol et al., 2010<sup>32</sup> | Level 1 |
| 25 | $O_{th}$ | 0.0510 | N/A | mg/L | threshold of oxygen concentration below which to switch from oxphos to glycolysis | Jiang et al., 1996<sup>37</sup>; McKeown et al., 2014<sup>37</sup> | Level 2 |

First, we aimed to approximate how much excess glucose is supplied by the vasculature recruited by glioma cells. Glioma vasculature provides the supply of necessary nutrients, including oxygen and glucose. A tumor’s ability to induce angiogenesis is a key and early step in tumorigenesis, representing a hallmark of cancer. Glucose and oxygen supply are directly proportional to the level of tissue vascularization; therefore, we leveraged the comparison of vascular density in GBM sections of variable cell density to infer *g*_*excess*_ (see Methods). We approximate the relation between cell density per voxel *N*_*i*_ and *g*_*excess*_ *(i)* with a linear regression model:

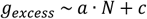

giving us an *R*^*2*^ = 0.59 (*P* = 0.00011; **Figure 2**(C). S3MB simulations qualitatively recapitulate this relation when considering vascular and cellular densities across multiple voxels (*R*^*2*^ = 0.321, P = 0.003; Supplementary Figure 11).

While digital pathology allows for an approximation of resource access in brain regions dominated by GBM cells, it does not inform about resource access in the surrounding brain parenchyma. The resources that glioma cells have access to as they invade from the tumor core into the surrounding brain vary considerably and glucose gradients influence cell migration (Supplementary Figure 7D). Moreover, cerebral perfusion varies more than 2-fold across various brain regions^55^ (e.g. vermis vs occipital lobes). Together these findings suggest that a brain-scale quantification of resources can improve our predictive power for disease progression.

To approximate resource distribution in the human brain we analyzed three public medical imaging datasets (**Figure 2**E), to derive an atlas of glucose uptake (**Figure 2**F), oxygenation (**Figure 2**G) and stiffness (**Figure 2**H) of the human brain (see Methods). We then asked whether age influences the spatial distribution described by the three atlases. The human brain, including its metabolic composition and physical properties, has been shown to change with age. For example, brain glucose levels increase with age^11^, while cerebral perfusion decreases with age at a rate of 0.45% per year^56^. Brain stiffness varies 2-fold between white and gray matter tracts and decreases with age^12^. We calculated the Pearson correlation coefficient between brain tissue stiffness and patient age across voxels and identified 11,853 (0.63%) and 535,205 (29%) voxels for which stiffness increased and decreased with age respectively (Supplementary Figure 6; FDR corrected p-value<0.05). The same analysis was also performed for glucose and oxygen (Supplementary Figure 6B,C), however we did not find significant associations with age after p-value correction, possibly due to the reduced number of subjects and biased age distribution. For glucose, there was a regional enrichment of either high-or low correlation coefficients in certain brain regions, suggesting significance would be reached if considering a lower spatial resolution. Our results confirm previous reports^11,12,57^ showing that age significantly affects resource distribution in the brain.

### S3MB can recapitulate regrowth of a primary GBM after the first surgery

Previous sections have shown how analysis of imaging data, previous modeling efforts, and clinical studies^58^ can quantify dependencies between resource availability in the TME and tumor cell phenotypes (in the form of ranges for parameter values listed in Table 1). Here we leverage clinical and genomic data obtained from a primary GBM patient to further narrow down these ranges for a small subset of these parameters, in particular those governing how cells respond to variable Glucose levels (Table 1), by either migrating (*V*_*max*_), proliferating (α_g_) or dying (α_d_, δ_R_).

We performed whole genome sequencing (WGS) of four surgical specimens collected during the 1^st^ and 2^nd^ surgeries of the patient. We used HATCHet (Hollistic Allele-specific Tumor Copy-number Heterogeneity)^59^, an algorithm that infers clone- and allele-specific somatic copy number alterations (SCNAs) jointly across multiple tumor samples, to quantify the clonal composition of the GBM and how it changes between the two surgeries (Supplementary section 4.1). Hereby each clone is defined by a unique set of SCNAs (Figure *3*A). Hatchet identified two aneuploid clones of ploidy 1.98 and 2.29 respectively (further referred to as low- and high ploidy clone). The low ploidy clone was dominant at the time of the first surgery and became even more dominant upon recurrence (**Figure 3**A). MRI images (T1 POST, FLAIR) were available before and after each surgery. T1 post and T2 flair scans from all four timepoints were linearly registered, followed by nonlinear transformation to MNI152 space. Enhancing and tumor infiltrating regions were segmented from T1 post and T2 flair scans respectively (**Figure 3**B). This data was used for in-silico model fitting of four model parameters (**Figure 3**C-J): (i) maximum migration speed (*V*_*max*_); (ii) glucose dependent death constants of the two aneuploid clones (β_*d, i*_ in eq.), (iii) glucose dependent growth constants of each clone (β_*d, i*_ in eq. *s*5) and (iv) starvation shrinkage rate (δ_R_ – how effective lack of resources is in killing tumor cells). Hereby the relative differences in glucose dependent migration between low- and high ploidy cells were retained as inferred from in-vitro experiments (shown in Supplementary Figure 7), with *V*_*max*_ scaling them to the in-vivo setting. Parameters (i - iv) were fit to the data in two iterations, one representing the average tumor cell population, the other distinguishing between the two clones (see Supplementary section 6). All other parameters were fixed as shown in Table 1.

**Figure 3:**
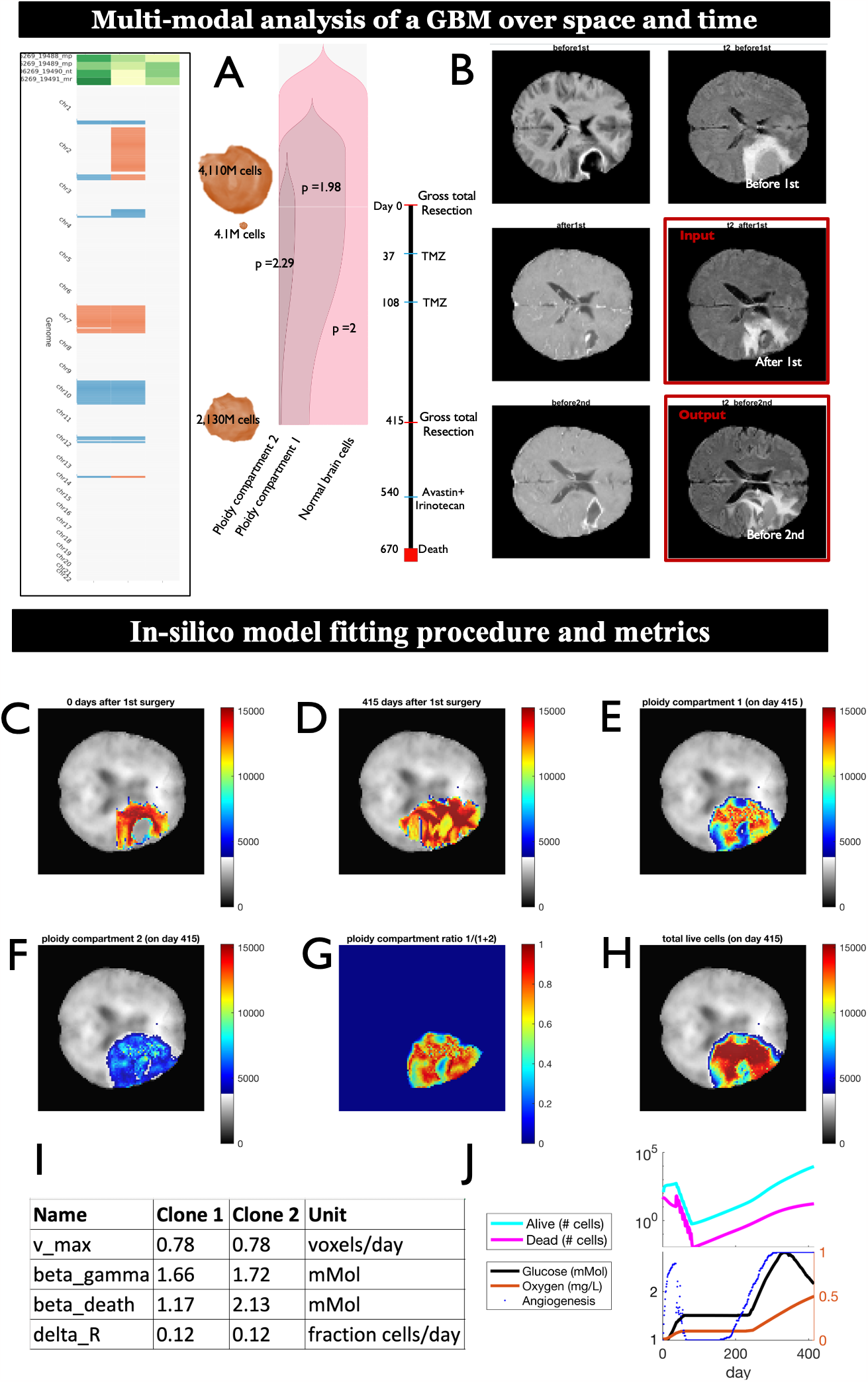
Multi-modal analysis of a GBM over space and time to inform mathematical modeling. (A) Co-clustering of clones (rows) detected with HATCHet at 2 different timepoints, shows changes in SCNAs (columns). (B) Genetic changes in primary and recurrent GBM clones shown relative to clinical events. 3D segmentation of T1 POST estimates tumor cell counts before and after the first and second surgery. (C,D) MRI-derived tumor cell densities recorded at 0 days (C) and 415 days (D) after the first surgery. (E-H) spatial tumor cell densities simulated by S3MB 415 days after the first surgery are shown for the low-ploidy subpopulation (E), the high ploidy subpopulation (F), the fraction of the low-ploidy subpopulation (G) and the two subpopulations in aggregate (H). Simulation used the tumor cell densities shown in (C) and subpopulation composition shown in (A) as initial condition. Background shows brain stiffness atlas used in the simulations. (I) parameters used for the simulation shown in (C-H). Parameters were optimized to recapitulate tumor cell densities shown in (D) and subpopulation composition shown in (A). (J) Temporal dynamics of angiogenesis, oxygen, glucose, dead cells and live cells. Sudden drop in glucose reflects dominance of glycolysis, until vasculature restores oxygen levels to facilitate switch back from glycolysis to oxidative phosphorylation.

Simulation results suggest that both clones have similar glucose-dependent growth constants (Clone 2: 1.72 mMol and Clone 1: 1.66 mMol), but that the high ploidy clone (Clone 2) had a higher glucose-dependent death constant (2.13 vs. 1.17 mMol for Clone 1; **Figure 3**E,F,I). These differences, in conjunction with the regional glucose, oxygen and stiffness resulted in spatial differences in the distribution of the two clones, with the low ploidy clone dominating in the interior and the high-ploidy clone dominating at the outer edges of the tumor (**Figure 3**G). Summing up simulated cell representations of the two clones per voxel (**Figure 3**H), resulted in cell densities that approximately recapitulate cumulative T1 post and T2 flair scans recorded 415 days after the first surgery (**Figure 3**D). A look at the temporal dynamics revealed that adjuvant therapy (consisting of chemotherapy and radiation therapy with TMZ) effectively killed tumor cells. However, the tumor partially repopulates the resection cavity after the last cycle of radiation therapy, with large numbers of cells invading beyond the cavity into the surrounding brain parenchyma after day ∼140 (**Figure 3**J).

Together these preliminary results demonstrate the potential of S3MB to recapitulate re-growth of a primary tumor into the patient’s recurrent tumor.

### Resource access during early pre-malignancy can shape ploidy composition and late-stage tumor progression

The outcome of the competition between WGD positive and negative cells is a branching point that sets glioma precursor cells onto differential paths of evolution^6^ and ultimately renders them sensitive to different drugs^5,7^. Several studies across a wide range of cancer types suggest that this branching point happens early in tumor evolution^1,4,60,61^. I.e. if a tetraploid subclone does expand, it tends to do so early, during the premalignant phase^1,4^. A potential explanation for this temporal contingency is that cell density during early stages of tumor evolution is still comparatively low and competition for nutrients is not yet so strong. Verena Koerber et al. reports that the founder cell of IDH^WT^ GBM emerges approximately 2-7 years prior to diagnosis^62^. This years-long premalignant phase may thus represent a window of opportunity for expansion of either diploid or tetraploid clones. Here we address the question whether the location of the first premalignant cells within the brain may influence the propensity of a tetraploid clone to expand and reach fixation, by virtue of intra-brain differences in resource supply.

We used S3MB to model growth of a hypothetical pre-malignant population of 1,000 cells, consisting of a diploid and tetraploid clone (50:50% frequency) in two different regions of the brain: one with low access to glucose (average of 1.22 mM; **Figure 4**A), the other with higher glucose access (average 2.36 mMol; **Figure 4**B). Hereby previously inferred differences in glucose dependent death-constants between the low- and high-ploidy clones (**Figure 3**I) were extrapolated to account for the increased difference in DNA content between diploid and tetraploid clones (2-fold difference in ploidy as opposed to just 1.1-fold difference between clones shown in **Figure 3**A). Same was done for glucose dependent growth-constants (Supplementary section 6.2). We let diploid and tetraploid clones compete for three years, seeding the initial population into a single voxel in each of the two different brain regions (**Figure 4**C,D). Its increased resistance to starvation induced cell death allowed the diploid clone to escape from the glucose poor brain region, whereas the tetraploid clone went extinct (maximum cell representation across all voxels fell below 1 cell on day 136). In contrast, when both populations were seeded into a glucose rich brain region, both clones still co-existed at the end of the simulation (**Figure 4**A,B). Hereby expansion of the high ploidy clone was dependent on activation of angiogenesis (**Figure 4**B).

**Figure 4:**
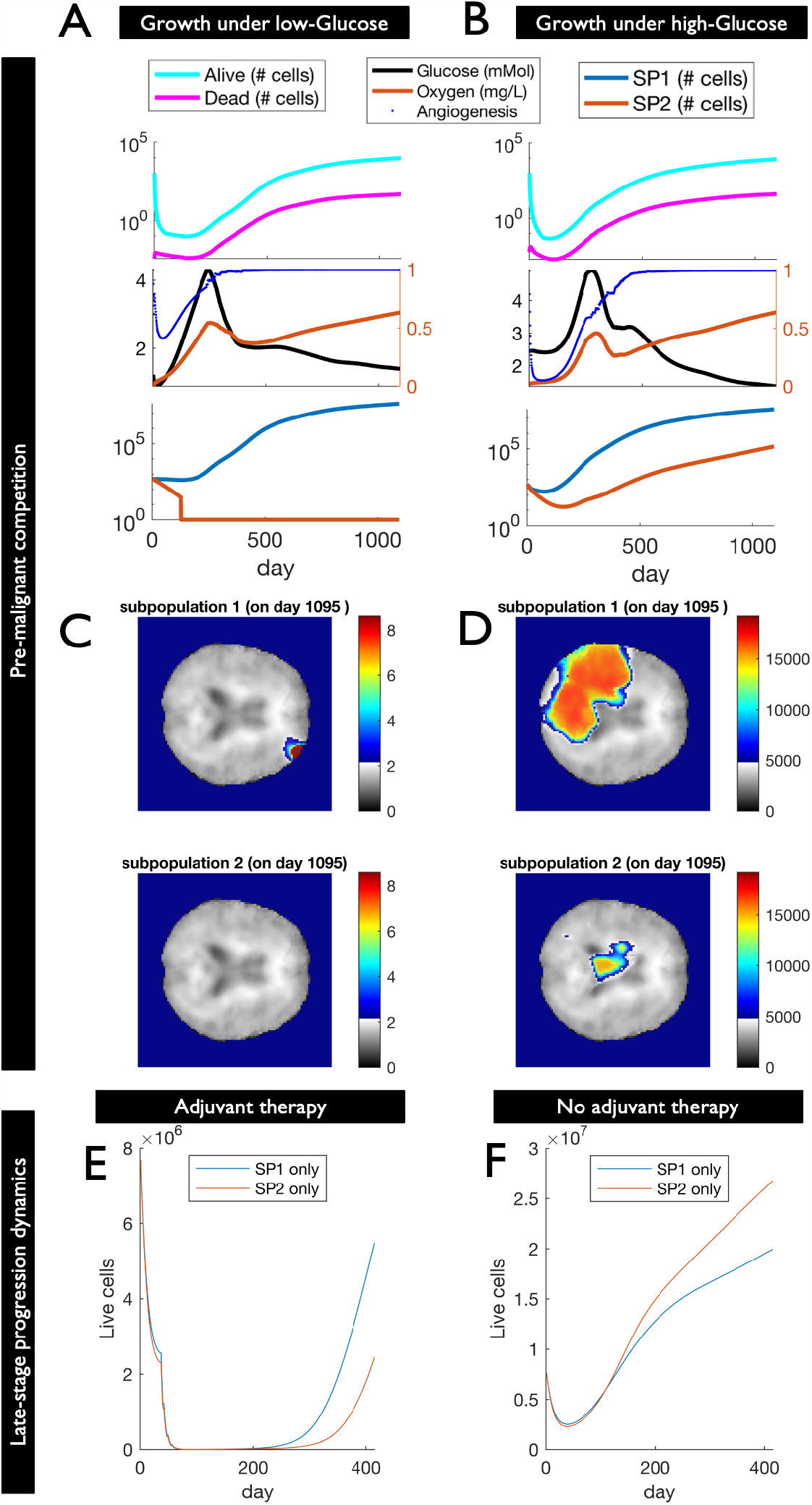
Resource access during early pre-malignancy can shape ploidy composition and late-stage tumor progression. (A-B) Growth of a hypothetical pre-malignant population of 1,000 cells, consisting of a diploid (50%: blue) and tetraploid clone (50%: red), was simulated in different regions of the brain: one with low access to Glucose (A), the other with higher Glucose access (B). Glucose-, Oxygen-, Angiogenesis, live and dead cell counts are shown alongside clonal composition. The tetraploid clone goes extinct in the low-Glucose area, but not in the high-Glucose brain region. (C-D) spatial distribution of the diploid (top) and tetraploid clones (bottom) is shown after 3 years for in-silico tumors emerging from low (C) or high Glucose brain regions (D). (E,F) Growth of a hypothetical GBM population consisting of 2.1M diploid (blue) or tetraploid cells (red) is simulated in the same brain region (as informed by post-surgery MRI in Figure 3D). The diploid GBM is predicted to progress and recur faster than the tetraploid GBM when exposed to adjuvant therapy (E). The reverse is true in the absence of therapy (F).

We next asked to what extent the outcome of this competition during early stages of tumor evolution affects tumor progression after detection. To address this question, we initialized S3MB with either a purely diploid or a purely tetraploid population using the same MRI-derived GBM cell densities shown in **Figure 3**C (recorded 0 days after the first surgery). When simulating chemo-and radiation therapy as indicated in the patient’s clinical record, the diploid tumor was predicted to recover from therapy faster, leading to an initially faster recurrence than if the population was purely composed of tetraploid cells (**Figure 4**E). When we repeated these simulations without any adjuvant therapy, the tetraploid tumor had a ∼2-fold higher rate of progression than the diploid one (**Figure 4**F). The relative fitness of purely diploid and tetraploid populations differs between the two scenarios (presence vs. absence of adjuvant therapy), because of the effect therapy has on the stiffness and resource supply of affected brain regions. Together these results support the hypothesis that resource access during tumor initiation can influence clonal interference between low- and high-ploidy clones, with implications for subsequent tumor evolution and recurrence.

Compared to intra-brain differences in resource supply, the differences observed between organs can be larger. For example, differences in oxygen concentration can span an order of magnitude^63^. If these differences between organs are consistent enough and if resource access does indeed influence the fixation rate of tetraploid cells, we would expect to see a correlation between the average concentration of a given resource per organ and the incidence of WGD in tumors originating in that organ. To test this hypothesis we compared median partial pressure of oxygen in human tissues^63,64,65^ to the median ploidy of cancer types that arise in the respective tissues (based on a previously published analysis of WGD in TCGA data^66^). We observed a significant correlation between these two variables (Spearman r = 0.65; P = 0.002), with the incidence of WGD being more than two times higher in organs with high oxygen partial pressure such as lung, compared to organs with low net oxygen (e.g. prostate gland; **Figure 5**A).

**Figure 5:**
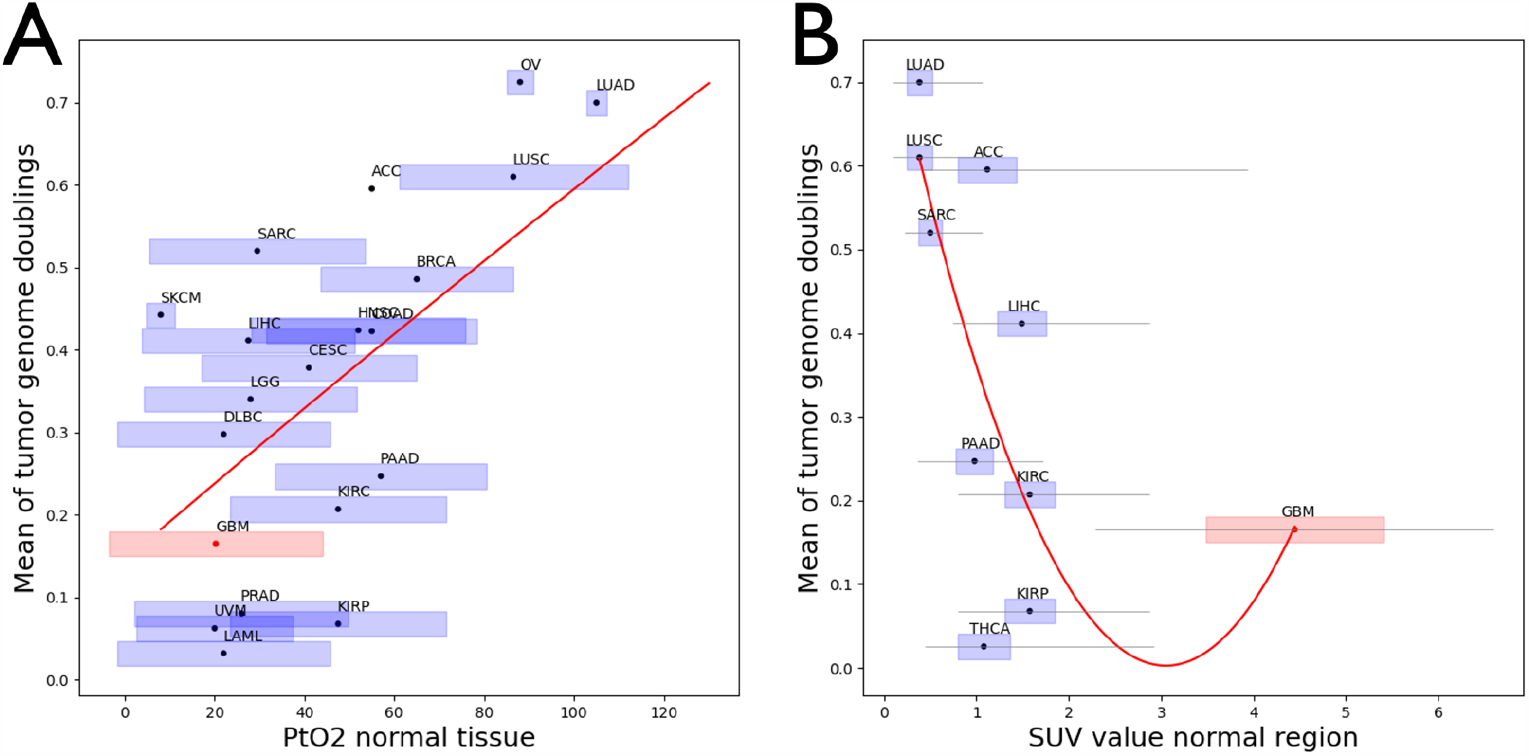
The average ploidy of 20 different cancer types is positively correlated with the oxygen levels recorded in their respective organ of origin (**A**; Spearman r = 0.65, P = 0.002) and negatively correlated to the glucose uptake rates in the organ of origin (**B**; Spearman r = -0.70, P = 0.026).

To quantify glucose limitations and how they relate to ploidy, we performed a multi-organ analysis of PET/CT scans from patients without underlying malignancy. Segmentation of 10 different organs from 397 PET/CT scans^67^ was performed with MOOSE^68^. We observed a negative correlation between organ-wide population-average estimates of glucose uptake and the average ploidy of cancers growing in the respective organ (**Figure 5**B). Assuming higher glucose uptake rates reflect fiercer competition for glucose, this result also suggests that lack of glucose imposes a higher fitness penalty on cells of higher ploidy.

Together these findings support the hypothesis that resource access shapes the evolution of ploidy, influencing the proliferation-, death- and migration rates of cells in a ploidy-dependent manner.

## Conclusions and perspectives

We present results from an integrated omics-, pathology and medical imaging analysis of primary and recurrent GBM. We use this multi-modal analysis to fit a Stochastic State-Space Model to recapitulate re-growth of enhancing primary tumor margins into the respective recurrent GBM detected in the same patient. The fitted model is then used to simulate growth of a mixed population of near-diploid and near-tetraploid cells at various locations of the brain, which differ in resource access.

We note a number of limitations of our approach. The first one is imposed by the unknown relevance of some of the literature-inferred parameters (Table 1) to the specific simulated setting. A critical next step will be to evaluate the sensitivity of S3MB to variability within the parameter ranges listed in Table 1. A second limitation is that differences in factors such as tissue pH or vessel wall thickness and luminal diameter were not taken into account when approximating excess resource supply in brain regions dominated by tumor cells. Blood vessels in GBM are known to be abnormal and highly permeable^69^, which is likely to further increase heterogeneity in resource supply and influence disease progression. Thirdly, we did not model differential sensitivity to radiation-therapy and TMZ between cells of different ploidy. While these differences likely exist^5,7^ and while S3MB offers the flexibility to incorporate them, we chose to neglect this aspect in favor of a focus on understanding resource allocation and how it pertains to DNA content. Lastly, it was not taken into account that ploidy makes a proportional impact on cell size, so each voxel can have less numbers of cells with high ploidy in comparison to near-diploid cancer cells. Despite these limitations, our approach offers a number of principled steps towards integration of a highly heterogeneous and multi-modal dataset into a mathematical model. This includes deriation of three atlases of resource distribution in the human brain, that can be overlaid with MRI scans from individual brain tumor patients. This resource is available to the research community and has potential to to inform predictive modeling of glioma cell invasion. We find supporting evidence for a strong effect of age on the distribution of these resources, suggesting that correction for age is likely to further improve accuracy of these atlas-derived approximations.

Future studies will focus on incorporating further aspects of cancer biology into the model. In particular, WGD+ cells are known to later lose parts of their tetraploid genome to converge on a highly favorable and commonly observed near-triploid karyotype^70–72^. WGD represents a more viable path toward this favorable near-triploid karyotype, as the resulting tetraploid intermediate can explore the genotype space more effectively in search for fitter cell states^4,73^. Aneuploid and polyploid cells are also most susceptible to mitotic catastrophe^74^ – which is often followed by necrosis. Necrosis is the most common form of death of TP53 mutated tetraploid cells after mitotic catastrophe^75^. Whether necrosis or apoptosis is the dominant form of cell death will in turn also shape the energetic environment of the forming tumor, as intra-cellular energy can only be recycled back to the environment after apoptotic, but not after necrotic cell death.

Our results provide proof-of-principle for inferring inter-patient variability in the conditions that dominated at the onset of a patient’s glioma formation and comparing them to the tumor’s respective aneuploidy status at the time of detection. They suggest that intra-brain differences in glucose access can suffice to influence whether a WGD+ cell population can co-exist with or even outcompete a diploid cell population (Figure 4). An organ-wide association study of cancer ploidy and resource access at host tissue sites supports this conclusion (Figure 5), suggesting that variability in resource access alone may be sufficient to explain divergent evolution of tumor cell ploidy. Future studies will investigate how much of the wide variability in ploidy within individual cancer types^66,76^, can be explained by variance in resource access at tumor initiation.

Another topic of interest and intense future studies is how age-induced changes in resource access shapes the evolution of ploidy. GBM remains uniformly lethal, with a median overall survival of 14-16 months, despite aggressive therapy. In people aged 65 and above however, prognosis is even more dismal, with median survival of less than 7 months. While comorbidities explain some of the increased age-related risk, age-related changes in the tumor microenvironment likely also contribute. The *outcome of the competition between near-diploid and near tetraploid pre-malignant cells may be a crucial branching point in the evolution to malignancy*, potentially influencing the speed of GBM recurrence and invasion. Whether higher ploidy predicts fast recurrence however may depend on the resource context in which the cells are growing, arguing for a classification of TMEs by their propensity to select for or against WGD. Understanding the selective pressures that shape ploidy at the onset of glioma formation, we expect, will contribute to mimicking their reverse therapeutically after tumor detection.

## Data Availability

S3MB is implemented in Matlab, and is available at https://github.com/cloneredesignlab/IMOworkshop2022.

## Acknowledgements

We thank Dr. Sandy Anderson for organizing the yearly IMO workshop and inspiring all its attendees to ralley together in a race of innovative ideas to bring mathematical and experimental oncology into the clinic. We also thank the jury who awarded our team the 1st price in the 10^th^ IMO workshop of 2022, themed: cancer communities.

