## Supplemental methods, figures and tables for "Interactions between ploidy and resource availability shape clonal interference at initiation and recurrence of glioblastoma"

### Supplementary Information

#### 1. S3MB: a stochastic state space model of the brain

Several computational models that simulate GBM growth as a function of the brain microenvironment, using reaction diffusion PDEs, have been previously developed[14,15]. We build upon this work to develop a new model whose assumptions in part overlap with those made by the previously developed proliferation-invasion-hypoxia–necrosis–angiogenesis (PIHNA) model of glioma growth[16]. The PIHNA model partitions the tumour cells into subpopulations based on histological characteristics such as normoxia, hypoxia, necrosis and angiogenesis, modeling interactions between them. What prior mathematical models of GBM invasion have been grappling with, however, are three challenges: the integration of multiple data types, the impact of intra-tumor heterogeneity and model evaluation in patients[15]. To address these three challenges, we have developed a Stochastic State-Space Model of the Brain (S3MB). Our model discretizes the brain into voxels, and the state of each voxel is defined by brain tissue stiffness, oxygen, glucose, vasculature, dead cells, migrating cells and proliferating cells of various ploidies, and treatment conditions such as radiotherapy and chemotherapy with temozolomide (TMZ). Well established Fokker-Planck partial differential equations govern the distribution of resources (oxygen, glucose) and cells across voxels[17]. However, we modified these equations as described below to simplify the model and rapidly track the temporal changes in these variables. First, we tracked the stiffness of the brain tissue within each voxel, as tissue stiffness in conjunction with available resources dictates the migration potential of cells[18]:

$$S = \overbrace{StiffIC \cdot c_0^{T_r^*}}^{\substack{radiation-induced\ edema \\ lowers\ stiffness}} \cdot \overbrace{\left(1 + c_1 \cdot \frac{N + N_d}{\sigma} \cdot H(N + N_d - 0.05 \cdot \sigma)\right)}^{\substack{tumor\ cell\ crowding \\ increases\ stiffness}} \quad \ldots s1$$

, where H is the Heaviside function, $StiffIC$ is the stiffness atlas of the healthy human brain (described in detail in Suplementary section 2.3), $\sigma$ is the available space within a voxel and $T_r^*$ is the cummulative radiation dose in a given region received up until time t: $T_r^* = \int_{t^*=0}^{t} T_r$. The first term mimics acute edema[77] in irradiated brain regions, which for the chosen parameter ($c_0 = 0.987$) lead to approximately 55% reduction of the initial brain tissue stiffness[78] by the end of a standard of care cumulative dose of $T_r^* = 60gy$, in line with prior reports[79]. Tumor cells can remodel the existing brain tissue and increase its overall stiffness, which for the chosen parameter ($c_1 = 6.68$) yields a maximum stiffness of approximately 22.5 kPa[18]. $N_d$ is the number of dead cells and $N$ is the total number of live cells across all subpopulations $i$: $N = \sum_i n_i$. Given local oxygen ($N_o$) and glucose ($N_g$) concentrations, changes in dead cells ($N_d$) and live cells of different ploidy ($n_i$) over time are described by:

$$\frac{\partial n_i}{\partial t} = \overbrace{\gamma_i n_i \left(1 - \frac{N+N_d}{\sigma}\right) \Gamma_i(N_o, N_g)}^{proliferation} - \overbrace{d_i(N_o, N_g, T_r, T_c) \cdot n_i}^{death} + \overbrace{K_i(N_o, N_g, S) \cdot \nabla n_i}^{migration} \quad \ldots s2,$$

$$\frac{dN_d}{dt} = \underbrace{\textstyle\sum_i d_i(N_o, N_g, T_r, T_c) \cdot n_i}_{death\ due\ to\ resource\ limitation/therapy} - \underbrace{c_d N_d}_{clearance\ of\ dead\ cells} \quad \ldots s3,$$

The first term on the right hand side (rhs) of equation $1a$ describes the logistic growth of these cells given resources ($N_o, N_g$) and available space ($\sigma$) within a voxel, the second term describes the death rate of cells due to resource limitations or therapy ($d_i(N_o, N_g, T_r, T_c)$), and the third term describes the migration of cells across neighboring voxels, driven by optimal resource utilization and brain tissue stiffness ($K_i(N_o, N_g, S)$)[80,81]. Both live and dead cells occupy the finite space available per voxel. Equation $1b$ describes the change in dead cells in a voxel and $c_d$ is the clearance rate of these dead cells via autophagy or other related mechanisms.

The dependence of cell growth and death on resources and treatment given at time t is given by:

$$\Gamma_i(N_o, N_g) = \overbrace{\left(\frac{N_o}{\alpha_{g,i}+N_o}\right) H(N_o - O_{th})}^{Oxygen\ is\ growth-limiting\ resource} + \overbrace{\left(\frac{N_g}{\beta_{g,i}+N_g}\right) H(O_{th} - N_o)}^{Glucose\ is\ growth-limiting\ resou□□ce} \quad \text{...s4}$$

and

$$d_i(N_o, N_g, T_r, T_c) = \overbrace{\delta_{R,i} \cdot \left[1 - \left(\frac{N_o}{\alpha_{d,i} + N_o}\right) H(N_o - O_{th}) - \left(\frac{N_g}{\beta_{d,i} + N_g}\right) H(O_{th} - N_o)\right]}^{\text{starvation−induced death}}$$
$$+ \underbrace{\delta_{T_c,i} \cdot \left(\frac{N_o}{\alpha_{Tchemo} + N_o}\right)}_{\text{chemo−induced death}} + \underbrace{H(O - O_{th}) \cdot f(\alpha, \beta, T_r) + H(O_{th} - O) \cdot f(\alpha, \beta, {}^{T_r}/_{r^*})}_{\text{radiation−induced death}} \quad ... s5$$

, with $\delta_R, \delta_T$ denoting the starvation- and treatment induced shrinkage rates respectively (how effective lack of resources / treatment is killing tumor cells), $T_c, T_r$ are chemo- and radiation-therapy doses applied in a given region in a time interval $dt$ and $\alpha_{Tchemo}$ is a Michaelis Menten constant of local oxygen availability which affects the effectiveness of chemotherapy with TMZ. The function $f(\alpha, \beta, T) = (1 - e^{(-\alpha T - \beta T^2)})$ is an empirically derived term for radiation-induced cell death.

$K_i(N_o, N_g, S)$ is a flux vector to the neighboring voxels, defined as:

$$K_i(N_o, N_g, S) = v_{max,i} r(N_o, N_g) \left[s(S) + \frac{d}{dxy} s(S)\right] \quad ... s6$$

, where, $v_{max,i}$ is the maximum rate of cell migration for cells with ploidy i, and $a_{mig,i}\ and\ b_{mig,i}$ describe the biphasic dependence of cell migration on glucose availability as:

$$r(N_g) = e^{-a_{mig} \cdot N_g} \left(1 - e^{-b_{mig} \cdot N_g}\right) \quad ... s7$$

$$s(S) = \mathrm{c}_2 \cdot S^2 + \mathrm{c}_3 \cdot S\text{The}$$
$$s(S) = \mathrm{c}_2 \cdot S^2 + \mathrm{c}_3 \cdot S$$

We assume that resource perfusion within the brain tissue under normal conditions is maintained at homeostasis via existing vasculature. However, the presence of rapidly growing tumor cells consumes resources at a rate faster than it can be replenished. At the same time, tumors can recruit additional vasculature to bring in additional resources. Under the simplification that each voxel is small enough that vasculature within that volume evenly distributes resources throughout the voxel volume, but large enough that diffusion of nutrients across voxel boundaries can be neglected (characteristic diffusion length of nutrient reaction-diffusion – ~100 µm – is much smaller than voxel dimensions: ~4 $mm^2$). Thus, changes in resources (oxygen and glucose individually) within a voxel are modeled as:

$$\frac{dN_o}{dt} = -\iota_o \cdot N_o N + \omega_o V \quad ... s8,$$

$$\frac{dN_g}{dt} = \overbrace{-\iota_g \cdot H(N_o - O_{th}) \cdot N_g N}^{\substack{Glucose\ consumption \\ when\ OxPhos\ dominates}} - \overbrace{\iota_{g2} \cdot H(O_{th} - N_o) \cdot N_g N}^{\substack{Glucose\ consumption \\ when\ Glycolysis\ dominates}} + \omega_g \cdot V \quad ... s9,$$

$$p(V) = p_0 H\left(N + N_d - \frac{\sigma}{4}\right) \cdot \sum_j \frac{N_j N_{d_j}}{\left(N_j + N_{d_j}\right)^2} \quad ... s10,$$

$$V = logical\ (T_r == 0\ \&\&\ V == 0\ \&\&\ rand < p(V)) \quad ... s11.$$

In equations $2a$,$2b$, $\omega_g, \omega_o$ are the additional resource generation rates by recruited vasculature and $\iota_o := \omega_o \cdot \frac{N}{\sigma}$, $\iota_g := \omega_g \cdot \frac{N}{\sigma}$ are the excess resource consumption rates by a voxel of tumor cells. Hereby $\iota_{g2} := 16 \cdot \iota_g$, i.e. cells consume more glucose when oxygen is low. Vasculature recruitment probability is given by equation 2c, whereby $j$ tracks adjacent voxels. The recruitment of new vasculature is promoted by a voxel being at certain capacity (1/4 shown here) and surrounded by voxels that are stressed due to comparable ratios of live and dead cells. Before triggering cell death, nutrient starvation elicits an autophagic response, which in turn has been shown to mediate pro-angiogenic effects via AMPK/Akt/mTOR signaling[84] [64]. The ROS-ER stress-autophagy pathway has also been shown to induce angiogenesis in and ischemic myocardium model[85]. Equation 2c therefor assume that comparable ratios of live and dead cells in a voxel maximize the probability of angiogenesis. As more dead cells accumulate the efficacy of recruiting vasculature is assumed to decrease. If on the other hand the voxel is dominated by alive cells we assume vascular recruitment is unnecessary as the area is sufficiently vascularized. The presence of recruited vasculature ($V$) is modeled as a binary value of 0 or 1. We assume radiation therapy will immediately kill all angiogenic cells within the treated voxels[86,87].

The model does have several floating parameters, particularly in terms describing resource and ploidy dependent cell growth ($\gamma_i, \Gamma_i(R)$ ) and death ($d_i(R,T)$) rates, as well as tissue stiffness and resource dependent cell migration ($K_i(R,S)$) rates for cells with different ploidy. An overview of all 25 model parameters is given in Table 1. For 13 of these parameters we found evidence from literature supporting a value or range typical for GBM, or we had in vitro data specific for GBM to inform them (Level 1). Another 12 of these parameters had approximations available for multiple other cancer types or other cell types, covering a wider range. In particular parameters representing resource availability and resource-driven phenotypic transitions were not well described and the motivation behind the selected values is described in the following section.

##### 1.1 Motivation for literature-based parameters related to resource-driven phenotypic change

$O_{th}$ – threshold of oxygen concentration below which a metabolic switch from oxidative phosphorylation (OXPHOS) to aerobic glycolysis occurs. The parameter was established based on threshold $O_2$ concentration (0.5%) for which the induction of expression level of hypoxia-inducible factor-1α and -1β (HIF-1α/β), key cellular regulator of metabolic switch from OXPHOS to glycolysis, is maximal, as measured in cultured HeLa cells[37];.

$C_d$ – clearance rate of dead cells via autophagy or related mechanisms. The range for this parameter (0.48-0.78) is motivated by the quantification of macrophage engulfment of target cells triggered to undergo apoptosis, necroptosis, or ferroptosis over 24 hours[28] and by the abundance of tumor-associated macrophages and microglia in the GBM tumor mass[29].

$\alpha_d$ – oxygen-dependent death constant per cell type ($O_2$ concentration at which cell survival is half-maximal) was selected based on the results of study evaluating the proliferation of different cell types under oxygen deprivation *in vitro*[20]. Oxygen concentration as low as 0.5% did not alter the fibroblast or tumor cells survival, irrespective of HIF-1 status, while extreme hypoxia (<0.01%) led to a block in proliferation and a progressive loss of viable cells to 20%-30% at 72 hours, also irrespective of HIF-1 status. Values in range 0.0007-0.034 mg/L (0.01-0.5% at 36°C) are supported by other studies of tumor proliferation under extreme oxygen concentrations[37].

$\beta_d$ – glucose-dependent death constant per cell type (glucose concentration at which cell survival is half-maximal) was selected based on the results of a study examining impact of low-glucose conditions on the survival of lung cancer cells[21]. We digitized Figure 1 using WebPlotDigitizer[88] and arrived at parameter range between 0-2.9 mMol depending on the cell line and pH conditions. The impact of pH on parameter values was not evaluated in S3MB.

$\alpha_g$ – oxygen-dependent growth constant per cell type ($O_2$ concentration at which cell proliferation is half-maximal) was selected based on the results of study evaluating the proliferation of different cell types under oxygen deprivation *in vitro*[20]. Similar to $\alpha_d$, it was found that cells continue to proliferate with undiminished kinetics under 0.5% oxygen and slow down proliferation in lower $O_2$ concentrations, as reviewed by McKeown et al[37].; $\alpha_g$ was assumed to be slightly higher compared to $\alpha_d$.

$a_{Tchemo}$ – oxygen at which chemo-shrinkage rate is half-maximum was selected based on temozolomide IC50 fold difference under hypoxia vs normoxia[34].

$c_0$– radiation-induced change in stiffnes – was the parameter value that led to a reduction of initial brain tissue stiffness by 55%, consistent with radiation-induced oedema[25]. Specifically, reduction in brain tissue stiffness is assumed to happen incrementally after each radiation cycle, without significant temporal delay[77]. A 55% reduction is achieved upon completion of a standard of care radiation therapy regimen, consisting of 60 Gy administered over 30 cycles.

$c_1$– change in stiffness from remodelling by tumor cells – Given StiffMax, the maximum GBM tissue stiffness of 22.5 kPa[18] and StiffIC, the average stiffness of healthy brain as quantified in our MRE derived stiffness atlas, we assume: StiffIC *(1+c1) = StiffMax and solve for $c_1$.

#### 2. Deriving oxygen, glucose and stiffness atlases of the human brain

We derive glucose uptake, oxygen and brain tissue stiffness atlases as follows.

##### 2.1 Glucose level atlas

An atlas of glucose-corrected standardized uptake values (SUVs), which inversely correlates with regional glucose availability, is available from a public database of 37 healthy adult human brains. It is comprised of [$^{18}$F]FDG PET, T1 MRI, FLAIR MRI, and CT images for each brain[47]. The PET scans were already coregistered to their respective T1 anatomical scans. To allow us to create an average brain for the purpose of our model simulations, all brains needed to exist in a standard space in which they could be averaged voxel-wise; to complete this, the scans were aligned with the MNI ICBM 152 2009c Nonlinear Asymmetric template[89] space using a linear followed by a nonlinear transformation with the Analysis of Functional Neuroimaging (AFNI) function 3dQwarp. One aged stiffness scan had to be excluded because of poor registration.

Following co-registration and registration to MNI space, PET images were normalized to obtain SUV and Standard Uptake Value ratio (SUVr) images at a $\leq$ 5% risk of Type 1 error[47]. These are used here and stratified by the patient's age. To approximate glucose levels, we assumed SUVr values are inversely correlated to glucose uptake[48,49] and that glucose concentration in the brain ranges between 1.0-2.7 mMol[50]. This was achieved through reflection of SUVr values, followed by scaling to the known range of brain glucose concentrations: $N_g = scale(-SUVr, [1.0, 2.7\ mMol])$. Since SUVr values are dimensionless, the scaling also assigns a unit to the resulting estimate $N_g$.

##### 2.2 Oxygen level atlas

The quantitative BOLD (qBOLD) data from seven volunteers was downloaded from the Oxford Research Archive site[51]. All image processing was performed with Quantiphyse[52]. To improve signal-to-noise ratio, four slices of 1.25mm thickness had been averaged together and there was slice gap of 2.5mm between resulting 5mm slices. We modified the NIFTI header to correctly report the actual slice interval of 7.5 mm, which allowed the qBOLD scans to align well with the anatomical scans. A linear registration and motion correction available in Quantiphyse[90], was used to correct any motion defects between timepoints of the qBOLD data. To suppress isolated noisy voxels we performed sub-voxel smoothing with a smoothing kernel of 1.5 mm. The mean reversible transverse relaxation rate (T2), the oxygen extraction fraction (oef) and the deoxygenated blood volume (dbv) were inferred with Bayesian modelling using the qBOLD widget in Quantiphyse[53]. The method deconvolutes three major confounding effects of qBOLD: (i) Cerebral Spinal Fluid (CSF) signal contamination; ii) Macroscopic magnetic field inhomogeneity and iii) Underlying transverse T2 signal decay[53]. To register all data to the Montreal Neurological Institute coordinate system (MNI) a DEEDS deformable registration[91] was performed to the anatomical images and the registration matrix was then applied to all the qBOLD data. Finally, an atlas of dissolved oxygen concentration (dO2) was derived from dbv and oef, in units of mg/L as:

$$\mathrm{dO2} = \mathrm{dbv} * (1 - \mathrm{oef}) * \mathrm{O2cap}$$

, where $O2cap = 0.4655$ mg/L is the typical average oxygen partial pressure expected in capillaries[54]. One scan had to be excluded because of poor registration.

#### 2.3 Brain tissue stiffness atlas

Magnetic Resonance Elastography (MRE) scans were obtained from a database of 12+12 aged and young brains generated at the University of Edinburgh, UK[12]. The T1 scans were first skull stripped with AFNI's 3dSkullStrip function. Next, because the MRE scans only covered a portion of the brain, a pad of 15 zeros was applied to the edge of the scan using the 3dZeropad function. The MRE scan was then co-registered to the T1 scan via rigid body transformation with the align_epi_anat.py functions, using the "-giant_move" option and NMI cost function; for those subjects who failed alignment, the "-ginormous_move" option (3 brains) or eliminating the move option (one brain) was used instead. The MRE scans were then transformed to MNI space by the same protocol as the PET scans.

#### 2.4 Correlation with age

MRI scan demographics for age and sex are tabulated below in Table S1.

Table S1: MRI Scan Demographics

| | PET | MRE | Qbold |
|---|---|---|---|
| Number | 37 | 23 | 7 |
| Age, mean (range) | 35 (23 – 65) | 46.3 (19 – 73) | 27.9 (24 – 32) |
| Sex, Female n (%) | 20 (54%) | 12 (52%) | 4 (57%) |

We first considered mean voxel value within the white matter (WM), grey matter (GM), and the whole brain parenchyma (WM+GM). We utilized the FSL FAST automated segmentation tool to create masks of the WM, GM, and cerebral spinal fluid for all of the t1 scans associated with patients who received PET and MRE scans. We segmented with FAST options: 3 classes, main MRF parameter of 0.1, 4 iterations for bias field removal, and bias field smoothing of 20.0 mm FWHM. We masked the various functional scans with the segmentations using afni's 3dcalc function and and computed the voxelwise mean (excluding zero values) with afni's 3dBrickStats.

Because the coregistered MRE scans were somewhat distorted on the edges, incorporating values between 0 and the minimum MRE value, we thresholded the t1-aligned MRE scans with the minimum original MRE value. This prevented the incorporation of low value but non-zero voxels in the grey matter average calculation. The rest of the steps were the same as those for the PET scans.

We then proceeded to compute Pearson correlation coefficients between patient age and brain tissue stiffness, glucose, and oxygen in a spatially dependent manner to determine which brain regions were most significantly altered during aging.

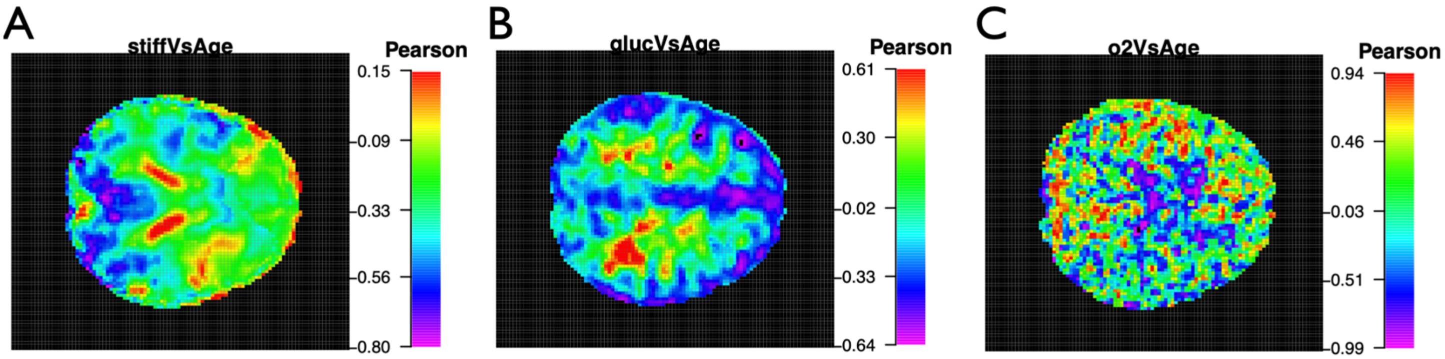


*Supplementary Figure 6: Map of Pearson correlation coefficients between patient age and brain tissue stiffness (A), Glucose (B) and Oxygen (C) across voxels.*

#### 2.5 Modeling the effect of surgery on oxygen, glucose and stiffness atlas

Because stiffness, glucose, and oxygen maps all share the same MNI coordinate system as the tumor cell density maps (see Supplementary section 4.3), we can model the effect of surgery on these environmental factors. Let $CAV$ denote the resection cavity map after a given surgery and $S, N_g, N_o$ are the stiffness, glucose, and oxygen maps. We assume tumor resection decreases stiffness as well as perfusion in the surgery-affected brain region (due to lack of a fractal pattern in blood supply provided by vasculature), as follows:

$$\mathrm{S} = \mathrm{S} - \frac{\mathrm{S}}{10} \cdot \mathrm{CAV}$$

$$N_g = N_g - \frac{N_g}{2} \cdot \mathrm{CAV}$$

$$N_o = N_o - \frac{N_o}{2} \cdot \mathrm{CAV}$$

#### 3. In-vitro experiments

##### 3.1 Cells

Experiments were performed using the near-diploid (2N) SUM-259 cell line. Near-diploid cells have previously been fused to derive a near-tetraploid (4N) lineage of the cell line as described in Miroshnychenko & Baratchart *et al.*[92]. All fused (4N) cells contained both, a fluorescent Mcherry staining to mark the nucleus and the GFP fluourescent probe to mark the cytoplasm.

##### 3.2 Incucyte transwell assay

DMEM media was prepared from powder (Sigma-Aldrich, 0453-0020) to vary glucose and dFBS concentrations (0.25-25 mM and 0.1-10% respectively). Media was filtered through a sterile 0.2mm filter bottle (Nalgene, 450-0020). Cells were starved for 12-24hrs before testing by replacing media with DMEM containing 1mM glucose and 1% dFBS. The Clearview Cell Migration plate (Sartorius, 4582) was used, if necessary, with a coating protocol with 25mg/mL collagen 1 (Corning, 354236) for an hour at 37ºC.  Cells were seeded into the upper chamber at 1,000 cells per well in 60mL DMEM with the appropriate glucose and dFBS concentrations. The lower portion of the wells were all filled with 200mL DMEM with 25mM glucose and 10% dFBS. The plate was imaged using the Chemotaxis assay in a Sartorius IncuCyte S3 at 37ºC & 5% $CO_2$ using 10x magnification, and images were taken every 2h for 48h.

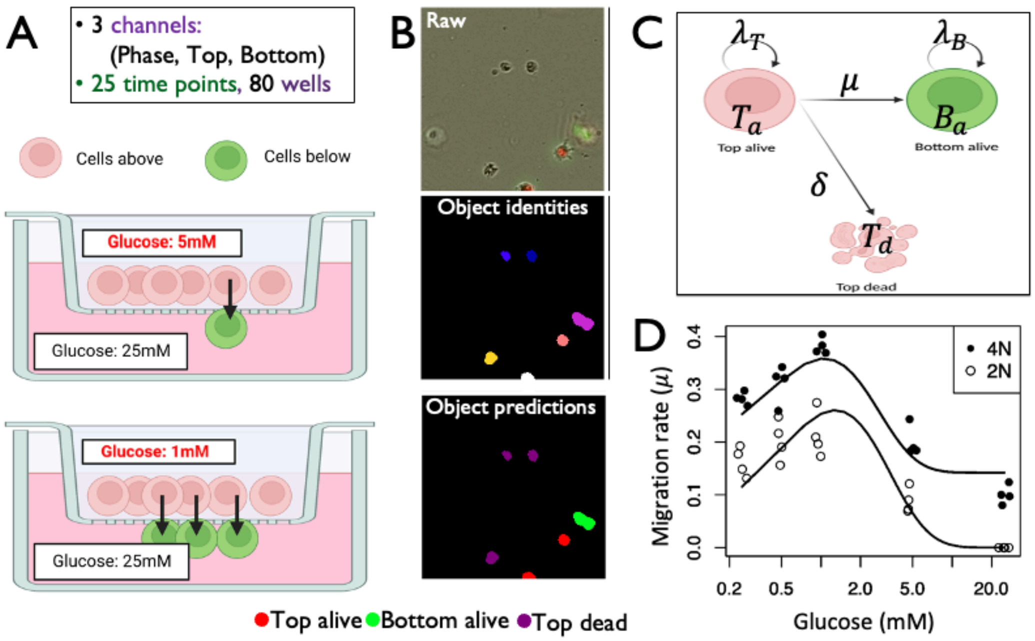


***Supplementary Figure 7: In-vitro quantification of cell migration in response to glucose deprivation****. (A) Overview of the chemotaxis trans-well assay. Cells were seeded in the top chamber under various glucose concentrations. Bottom chamber contained 10% FBS and 25mM glucose. Cells respond to the various gradients between top and bottom chamber, migrating through pores to reach the richer growth media at the bottom. (****B****) Images were acquired every 2 hours for 2 days, cells were segmented and classified according to their location (top vs. bottom) and cell state (live vs. dead). (****C****) Data was modeled using an ODE consisting of 3 compartments: bottom live, top live and top dead cells. Cells can migrate from top to bottom or die at various rates (μ and δ respectively). (****D****) Model shown in (C) was fitted to the data shown in (****B****) and the hereby inferred migration rates are plotted against the corresponding Glucose concentration on the top well. Datapoints are stratified by cell type (color-code).*

##### 3.3 Image analysis

We processed 6000 raw images from the Incucyte transwell assay (80 wells; 25 times points per well; and three channels Phase, Top, and Bottom); for this, we colored the top and bottom channels with red and green respectively, then we merged them and stacked the time points images associated with the same well, thus obtaining 80 files equal to the number of wells (80 wells).

After manipulating the raw images, the newly processed images were used for the creation of the training and test datasets. First, we selected only one frame (time point) per well sequentially, so all different times, from 0 to 24, were sampled throughout the files (wells) at least once. Second, we cropped a random region of 200 × 200px from each frame selected. All this process was done twice to create both training and test datasets. Each dataset was then composed of 80 frames.

Succeeding the data preparation, we used the image classification and segmentation software *ilastik*[44] to obtain cell counts and cells' spatial information.  We trained the model (using the training dataset) by first assessing a pixel classification to separate cells from the background. Then, we performed the object classification task. In this last part, cells were classified manually into three labels: top alive, bottom alive, and top dead. We identified top and bottom alive cells with the help of the fluorescent staining - red and green colors we used previously-; meanwhile, we recognized top dead cells by eye inspection and by comparison with the changes observed in the time evolution of the original files for each well. Finally, the classification was validated using the test dataset; for this, we compared the classifier's cell counts output for the test dataset by manually counting the number of cells in each frame of the dataset.

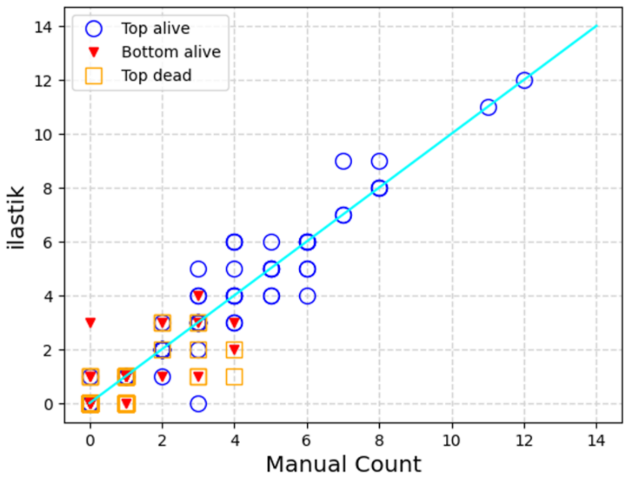


***Supplementary Figure 8: Validation of cell counts derived from the chemotaxis Sartorius IncuCyte assay**. We processed 6000 raw images from the Incucyte transwell assay (80 wells; 25 times points per well; and three channels Phase, Top, and Bottom). To create a training and test dataset, we cropped two random regions of 200×200px from each frame selected. We used the image classification and segmentation software ilastik28 to classify cells as "top alive", "bottom alive", and "top dead". Shown here is the comparison between pipeline's output on the test dataset (y-axis) and manual counts (x-axis) per each of the three classes.*

##### 3.4 ODE model of cell migration in response to glucose deprivation

In order to fit the data (cell counts) obtained from the image analysis described in section 3.3, we developed an ODE model which describes the temporal evolution of three cell compartments: Top alive ($T_a$), Bottom alive ($B_a$), and Top dead, ($T_d$), as follows:

$$\frac{dT_a}{dt} = \lambda_T(G_T)T_a - \delta(G_T)T_a - \mu(G_T)T_a$$
$$\frac{dB_a}{dt} = \lambda_B B_a + \mu(G_T)T_a$$
$$\frac{dT_d}{dt} = \delta(G_T)T_a$$

In this model, $T_a$ cells can proliferate at a rate $\lambda_T$, migrate to the bottom chamber at a rate $\mu$ and become $B_a$cells, or die at a rate $\delta$ and become $T_d$ cells. Furthermore, since there are enough resources at the bottom chamber, cells at the bottom ($B_a$) can only proliferate at a rate $\lambda_B$ and do not die. Additionally, all parameters were considered to depend on the glucose concentration in the top chamber $G_T$, but only the proliferation rate of bottom cells $\lambda_B$ remained constant. The reason for keeping $\lambda_B$ constant is that the conditions at the bottom chamber are always the same among wells (10% FBS and 25 mM of glucose), the only difference is the various glucose concentrations at the top chamber.

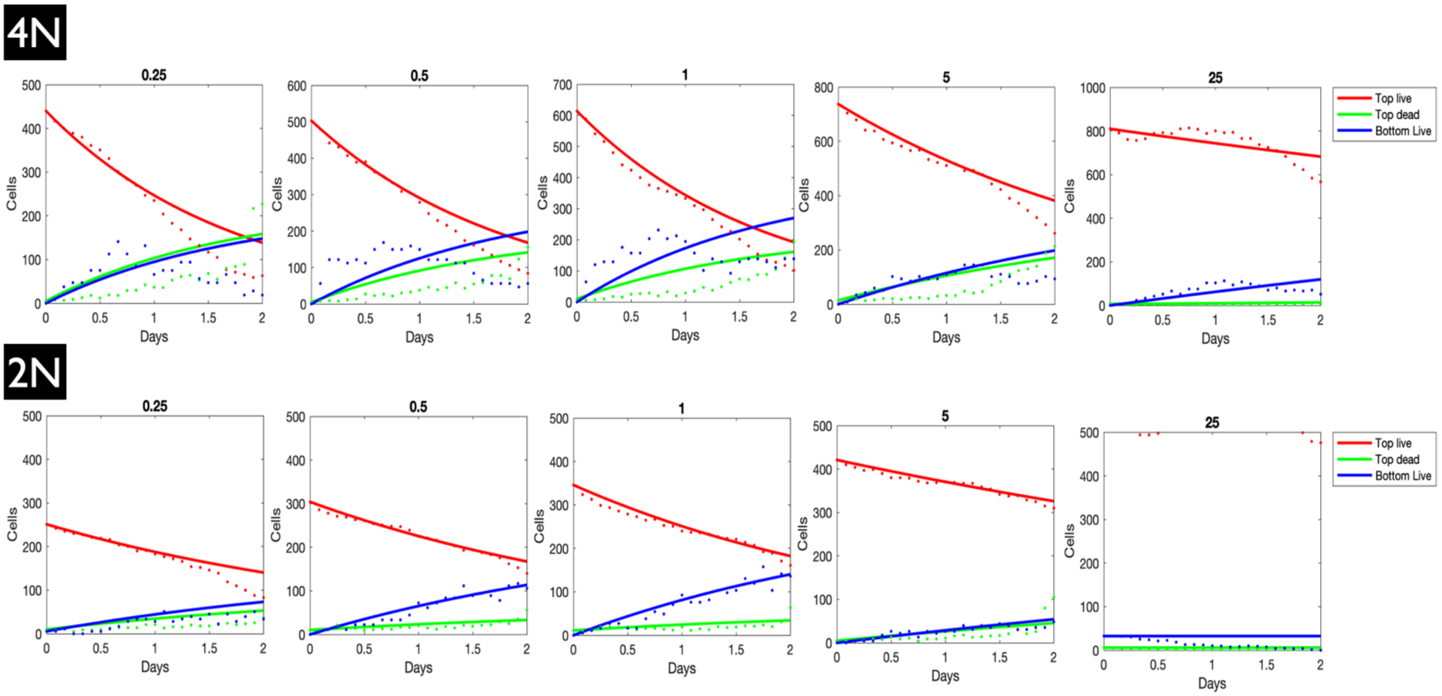


***Supplementary Figure 9: Fitting model of cell migration in response to Glucose deprivation to experimental data.*** *We developed an ODE model of cell migration in response to Glucose deprivation, which describes the temporal evolution of three cell compartments: Top alive ($T_a$), Bottom alive ($B_a$), and Top dead, ($T_d$). In this model, $T_a$ cells can proliferate at a rate $\lambda_T$, migrate to the bottom chamber at a rate $\mu$ and become $B_a$cells, or die at a rate $\delta$ and become $T_d$ cells. To infer how glucose concentration in the top chamber affects migration, proliferation and death rate of $T_a$ cells, we fitted the ODE model (curves) to the data (points) from the five top chamber glucose concentrations (shown here on top of each panel: 0.1, 0.5, 1, 5 and 25 mM). Fitting results are shown for one (out of four) representative replicate.*

We fitted the ODE model to the data from the five top chamber glucose concentrations (0.1, 0.5, 1, 5, and 25 mM) simultaneously. To do so, we used the *lmfit* library in Python and the method of Levenberg-Mawquardt. In total, 16 parameters and 15 initial conditions were allowed to vary during the fitting.

#### 4. Multi-modal analysis of a GBM over space and time

##### 4.1 Sequencing

We performed whole genome sequencing (WGS) of four surgical specimens collected during the 1$^{st}$ and 2$^{nd}$ surgeries of a primary GBM patient. Sequencing adapters were trimmed from whole genome sequence data using CutAdapt[93]. Trimmed sequences were aligned to the GRCh38 human reference using the Burrows-Wheeler Aligner (bwa mem)[94] . Duplicate identification, insertion/deletion realignment, quality score recalibration, and BAM sorting and indexing were performed with PICARD, the Genome Analysis ToolKit (GATK)[95], and samtools[96]. Germline variants were identified using GATK UnifiedGenotyper on all tumor and normal samples simultaneously. Allele frequencies for all tumor samples were extracted at normal sample heterozygous variant positions from the GATK output VCF file using custom Perl scripts. Copy-number based subclonal architecture and how it changes between the two surgeries was derived using HATCHet[59]. Briefly, read depth was counted in 50kb bins. Read depth and B-allele frequency were integrated in the 50kb bins, which were then prepared for HATCHet analysis. The clustering step (cluBB) used tolerancebaf=0.08 and tolerancerdr=0.2, and clusters were evaluated visually for appropriate cluster assignment. HATCHet was run with a range of 2 to 8 clones, minimum clone proportion of 0.03, ghostproportion of 0.35, and upper bound on relative increase of objective function of 0.6.

##### 4.2 H&E image analysis

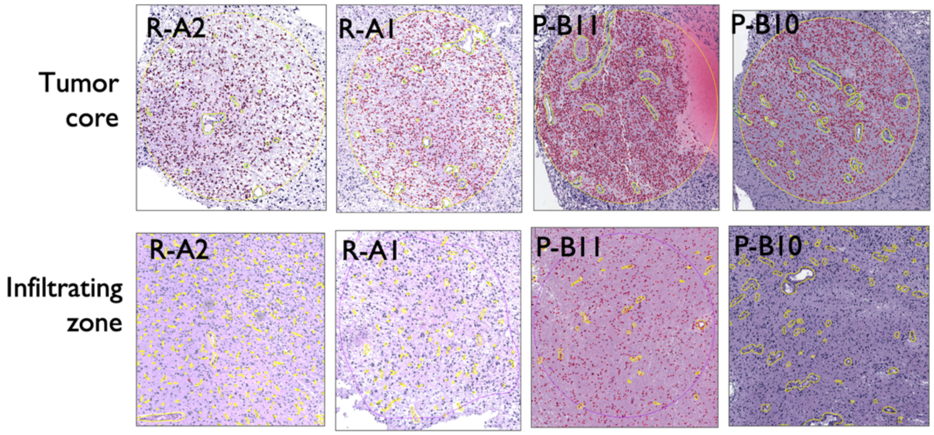


*Supplementary Figure 10: Vasculature in GBM core and adjacent normal tissue.*

Goals of the H&E image analysis were to: (a) estimate carrying capacity as the maximum cellularity per voxel and (b) estimate $\omega_g, \omega_o$ as the additional Glucose and Oxygen generation rates by recruited vasculature. We achieved this in four steps.

In the first step, four samples of GBM H&E stainings were analyzed by a histopathologist. Multiple tumor core and infiltrating zones were identified in each slide as circular regions of interest (ROI) of 1 mm diameter each. The infiltrating zone (IZ) was defined as a tumor region directly adjacent to the non-neoplastic tissue, whereas the tumor core (TC) was defined as a tumor region a few millimeters from the tumor/non-tumor border. Using the QuPath software[45], the vascular architecture of each ROI was analyzed, and the number of tumor cells was counted. The vascular architecture was approximated and marked manually, whereby differences in vessel wall thickness and luminal diameter were not considered. In turn, the number of tumor cells was counted automatically using algorithms applied in the QuPath software, where the basis for the measurement was the number of nuclei, their surface and shape. This yielded a total of 5 tumor core ROIs and 12 infiltrating zone ROIs, $ROI \coloneqq \{TC_1, \dots TC_5, IZ_1, \dots IZ_{12}\}$.

Because all ROIs have the same surface area, the total vascular area and cell count per ROI are proxies of vascular density and cell density respectively. We calculated $N$-- the cell density per voxel as:

$$N = n \cdot voxel_{size} / \overline{ROI_a}$$

, where $voxel_{size} = 4\, mm^2$, $n$ is the total number of cells detected per ROI and $\overline{ROI_a}$ is the average area of an ROI ($\sim 0.85 mm^2$). The carrying capacity was then calculated as $\sigma = max(N_x), x \in ROI$. The approximated value ($\sigma = 15{,}448\ cells/voxel$) was similar to estimates from literature[46].

Next, we calculated the total area occupied by vasculature ($V_a$) for each ROI. While both cell density and vascular density were higher in $TC$ regions than in $IZ$ regions, both variables changed in tandem in a linear fashion irrespective of region type (Supplementary Figure 11). We then estimated the additional glucose ($g_{excess}$) and oxygen ($o_{excess}$) recruited by tumor vasculature per each ROI, $i$, as:

$$g_{excess}(\mathrm{i}) = G_0 \cdot (\frac{V_{a,i}}{\overline{V_{a,IZ}}} - 1)$$

$$o_{excess}(\mathrm{i}) = \Theta_0 \cdot (\frac{V_{a,i}}{\overline{V_{a,IZ}}} - 1)$$

, where $G_0$, $\Theta_0$ are the average glucose (expressed in units of $mMol/voxel/day$) and oxygen ($mg/L/day$) generated per day by physiologic cerebral vascular supply. Above calculation makes the following assumptions:

a. Glucose/oxygen generation rate is directly proportional to the vascular density
b. As the vascular density is higher in regions dominated by tumor cells, the mean vascular density across all IZ ROIs approximates vascular density of normal brain regions. Thus, dividing by the average gives us the fold change of vascular density in tumor regions (and via $a$. also the fold change in glucose/oxygen generation rate).
c. Subtracting 1 from the outcome and multiplying by $G_0$ or $\Theta_0$ thus approximates excess glucose or oxygen generation rate by recruited vasculature.

d. $G_0 := \overline{N_g}$ is the average glucose recorded in the glucose atlas $N_g$ (see also Supplementary section 2.1).
e. $\Theta_0 := \overline{N_o}$ is the average oxygen recorded in the oxygen atlas $N_o$ (see also Supplementary section 2.2) .

As third step we then calculated $o_{excess}$ for all $TC$ regions $O_{excess} := \{i \in TC | \, o_{excess}(i)\}$ and approximated $\omega_o$ as the median excess oxygen produced by $TC$ regions: $\omega_o = \widetilde{O_{excess}} \cong 0.0026$. Analogue we calculate $G_{excess} := \{i \in TC | \, g_{excess}(i)\}$ and approximate $\omega_g$ as $\omega_g = \widetilde{G_{excess}} \cong 0.182$:

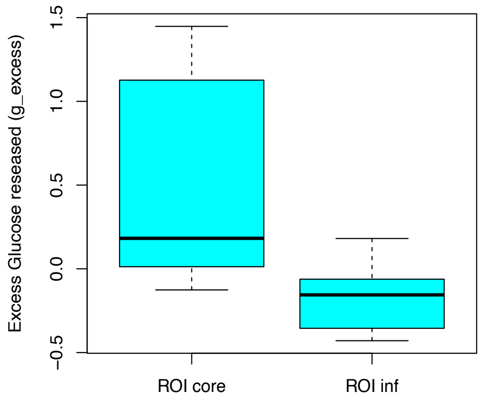


Finally, we approximate the relation between $N_i$ and $g_{excess}(i)$ across ROIs with a linear regression model:

$$g_{excess} \sim a \cdot N + c$$

giving us an $R^2 = 0.59$ ($P = 0.00011$; Supplementary Figure $11A$). Simulations qualitatively recapitulate this relation when considering vascular and cellular densities across multiple voxels ($R^2 = 0.321$, P = 0.003; Supplementary Figure $11B$).

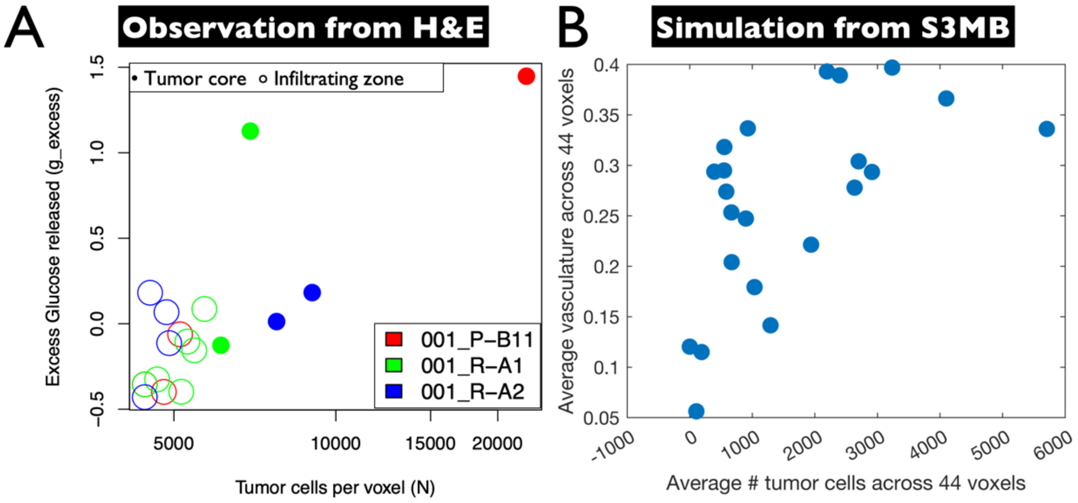


*Supplementary Figure 11*

#### 4.3 MRI analysis

MRI scans (T1 post, T2 flair) were available before ($B1, B2$) and after ($A1, A2$) two maximum safe resection surgeries. Masks were generated from the T1 scan, to cover the region occupied by solid tumor, and from the T2 flair to indicate the bounds of surrounding edema and infiltrative glioblastoma cells. The tumor masks were already coregistered to the T1 or T2 flair scans respectively, as the masks were generated from the anatomical scans. The anatomical scans and masks were first resampled to the cubic voxels that matched the smallest dimension of the voxels in the anatomical scans using the AFNI function 3dresample, to aid in registration to MNI space.

Next, the T1 scan (and its mask) was carried to MNI space using a linear followed by nonlinear transformation. The T2 flair scan was next coregistered to the T1 scan via an affine rigid body transformation and this transformation matrix was applied to the T2 mask. Both the T2 flair scan and the mask were then registered to MNI space using the same transformation as the T1 scan and mask. The tumors distorted normal cortical anatomy so the skull stripping was somewhat imperfect; a binary mask was generated from the standard MNI brains and applied to the MNI-registered brains with tumors to exclude residual skull for our simulations.

To approximate relative cell density (RCD) within the patients' brain from T1 post and T2 flair signals we used a previously described multiparametric regression model for mapping glioblastoma cellularity[97], giving us four relative cellularity matrices before and after 1st and 2nd surgery: $\mathrm{RCD_{B1}}$, $\mathrm{RCD_{B2}}, \mathrm{RCD_{A1}}, \mathrm{RCD_{A2}}$. To go from relative to absolute cell density (ACD), we assumed that the maximum RCD value before the 1st surgery corresponds to the tumor's carrying capacity (see Table 1), giving us a normalization factor:

$$f_{norm} = \frac{\sigma}{\max\,(\mathrm{RCD_{B1}})}$$

This factor was used to approximate ACD before and after each surgery as:

$$\mathrm{ACD_x} = f_{norm} \cdot \mathrm{RCD_x}$$

, where $\mathrm{x} \in \{\mathrm{B1, B2, A1, A2}\}$. A drawback of this approach is that T2 flair signal after surgery is not only influced by tumor cell density but also by inflammation caused by the surgery itself. To subtract the effect of inflammation, we enforced tumor cell density after a given surgery not to exceed tumor cell density prior to the surgery:

$$\mathrm{ACD_{A1}} = \min\,(\mathrm{ACD_{B1}}, \mathrm{ACD_{A1}})$$

$$\mathrm{ACD_{A2}} = \min\,(\mathrm{ACD_{B2}}, \mathrm{ACD_{A2}})$$

The underlying assumption is that surgery can only decrease, and not increase tumor cell density per voxel.

Lastly we set cell density inside the resection cavity to zero:

$$\mathrm{ACD_{A1}} = \mathrm{ACD_{A1}} - \mathrm{ACD_{A1}} \cdot CAV$$

where $CAV$ denotes the resection cavity map.

Because stiffness, glucose, and oxygen maps all share the same MNI coordinate system as the ACD maps derived above, we have access to proxies of these environmental factors throughout the tumor and the healthy brain tissue.

#### 5. Unit Tests

We performed a series of unit tests to verify the implementation of eqs. $s1$-$s11$. These tests verify the behavior of birth/death (Supplementary Figure 12A), diffusion (Supplementary Figure 12B), angiogenesis (Supplementary Figure 12C) and metabolic switches (Supplementary Figure 12D) in isolation from all other functionality. The exact conditions underlying each test and the respective expected outcome are summarized in Supplementary Figure 12.

A

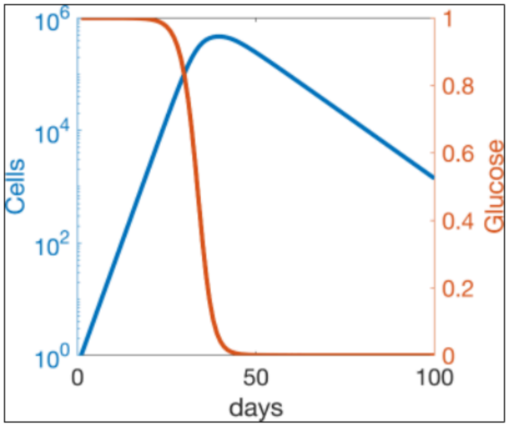


**Birth-Death Test**

- Optimal stiffness and uniform glucose concentrations
- No angiogenesis
- No migration
- Infinite carrying capacity
- Expected behavior:
  - **Cells grow at birth-deathrate until glucose is consumed**
  - **Cells die at deathrate once glucose falls below min**

B

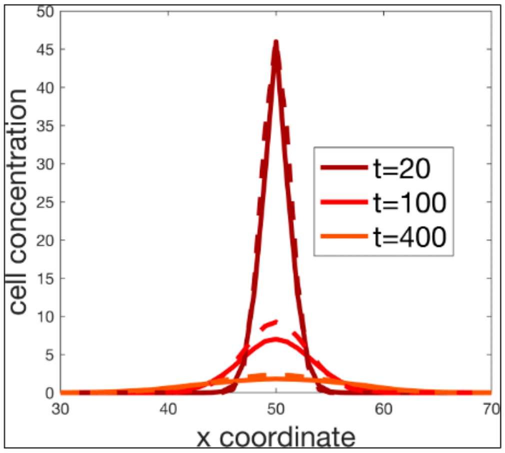


**Diffusion Test**

- Birthrate=Deathrate=0; **Migration rate >0**
- No angiogenesis
- Stiffness, resources: uniform throughout brain
- Cells seeded at high densities in a single voxel
- Expected behavior:
  - **Cell distribution agrees with solution to diffusion eq. in 2D from point source in infinite domain**

C

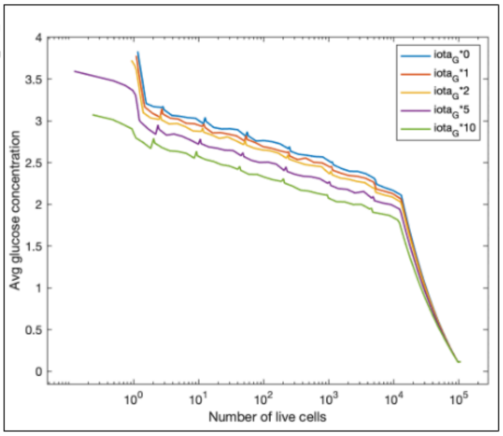


**Angiogenesis Test 1**

- Optimal stiffness and low Glucose concentrations (constant throughout the brain)
- No migration
- Cells seeded at 1/8 of their carrying capacity uniformly throughout the brain
- Expected behavior:

➢ **Total glucose concentration (free + consumed by cells) decreases with value of $iota_G$**

D

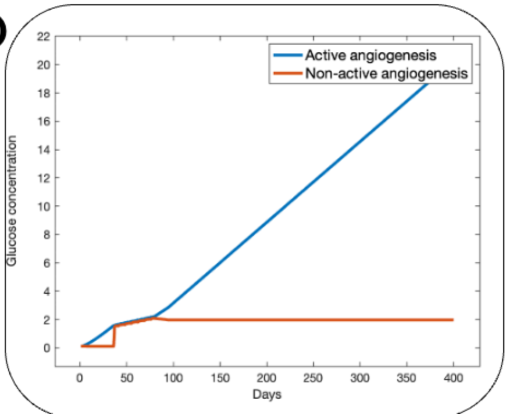


**Angiogenesis Test 2**

- Optimal stiffness and low Glucose concentrations (constant throughout the brain)
- No migration
- Cells seeded at 1/8 of their carrying capacity uniformly throughout the brain
- Expected behavior:

➢ **Voxels where angiogenesis is active have higher concentrations of Glucose (free + consumed by cells)**

*Supplementary Figure 12: S3MB unit tests. (A) Proliferation and apoptosis rates as a function of Glucose concentration. (B) Diffusion tests results. Comparison between theoretical predictions and simulations of cell distribution for a constant population with a positive migration rate. (C,D) Angiogenesis tests.*

For the diffusion test (Supplementary Figure 12B), we seed the initial cell population in a single voxel, precisely the middle one in position (50,50). We also keep constant population size by considering null birth and death rates. We don't consider angiogenesis and we set a positive migration rate for cells, together with constant resources of glucose and oxygen throughout the brain. We then plot the distribution of cells at different times of the cancer progression, and we compare these results with the following theoretical expectation (solution to diffusion equation in 2D from point source in infinite domani):

$$u(x,t) \approx \Delta x \Delta y \frac{u_0}{4\pi D(x,t)t} e^{-\frac{(x-x_0)^2}{4D(x,t)t}}$$

where:

- $\Delta x$ and $\Delta y$ are the distance between each voxel in the equispaced grid (in our case then they are equal to 1);
- $u_0$ and $x_0$ are respectively the size of population and the position of the voxel cells are initially seeded;
- $t$ is the number of days for which we are approximating the distribution;
- $D(x,t)$ is the approximation of diffusion from equation $s6$ for timepoint $t$, calculated as:

$$D(x,t) \approx \frac{1}{4} K(|N_o, N_g, S)$$

The simulations match the theoretical expectation for different time frames (Supplementary Figure 12B).

For the angiogenesis tests (Supplementary Figure 12C-E), we set cell migration to zero, and we seed the cells at 1/8 of the carrying capacity uniformly throughout the brain, together with low constant resources of glucose and optimal stiffness. We perform two different analyses for understanding the role of angiogenesis in tumor spreading and glucose concentration over time and space. In Supplementary Figure 12C, we show how the average glucose concentration over time and space is correlated to the parameters $\iota_G$. In the test in Supplementary Figure 12D, we perform an analysis of the correlation between glucose concentration and angiogenesis activity, showing that the first is much higher where angiogenesis is active. The glucose concentration on the y-axis is calculated by taking the mean over time among the voxels with active angiogenesis, more precisely:

$$g_a(t) = mean\left(N_g(t)|_{active\ vasc}\right)$$

$$g_{na}(t) = mean(N_g(t)|_{non-active\ vasc})$$

where $g_a(t)$ and $g_{na}(t)$ are respectively glucose concentration where angiogenesis is active and the one where angiogenesis is not active at time $t$.

#### 6. Data fitting

The S3MB domain was initiated from 4mm$^3$ voxels of the MNI space, with segmented T1 post and T2 flair regions encoding for different tumor cell densities per voxel. A representative slice was taken to include the lateral ventricles and used as the initial plane for the 2D simulations (**Figure 3**C), along with its WGS-derived ploidy composition (**Figure 3**A). We performed a parameter search to recapitulate the GBM's cell density and ploidy composition before the 2$^{nd}$ surgery. Fitting was done in two steps: (i) modeling just a single subpopulation and using tumor cell density alone for parameter optimization; (ii) modeling both subpopulations but fixing a subset of the parameters informed by step (i) to be identical for both populations, while the remaining parameters were further optimized. For (ii) tumor cell density as well as subpopulation composition were used for parameter optimization.

##### 6.1 Modeling a single subpopulation

Let $l$, L $\in \mathbb{R}_+^{n,n}$ be the predicted and observed spatial tumor cell density distribution (live + dead cells) on the day of the second surgery respectively (415 days after the first surgery). Then we define:

$$D_1 := \left\{ \left(1 - \frac{L_A}{l_A}\right)^2, \left(1 - \frac{l_B}{L_B}\right)^2 \right\}$$

, where $A \coloneqq \{i|\ before2nd_i < livedead_i\ \wedge\ before2nd_i + livedead_i > \varepsilon\ \}$ and $B \coloneqq \{i|\ before2nd_i > livedead_i\ \wedge\ before2nd_i + livedead_i > \varepsilon\}$. Hereby the role of $\varepsilon \coloneqq 0.05\sigma$ is to exclude true negatives (regions without tumor correctly predicted as such) from the cost function. This ensures smaller tumors will not have a better fit just because the brain contains more tumor-free regions. A global fitting procedure (MATLAB multistart) was used.

##### 6.2 Modeling both subpopulations

Let $p, \mathrm{P} \in \mathbb{R}_+^{1,2}$ be the predicted and measured proportions of the two tumor subpopulations respectively (see Figure *3*A), also on the day of the second surgery. We define:

$$D_2 := \left(1 - \frac{p}{P}\right)^2$$

The cost is then calculated as:

$$cost \coloneqq \sqrt{\overline{D_1} + \overline{D_2}}$$

In a first step, the following four parameters were fitted according to the approach described in 6.1: (i) maximum migration speed ($v_{max}$); (ii) glucose dependent death constant ($\beta_d$ in eq. $s5$ ), (iii) glucose dependent growth constant ($\beta_g$ in eq. $s5$) and (iv) starvation shrinkage rate ($\delta_R$). Then, two of these parameters ($\beta_d$ and $\beta_g$) were centered according to the results from the first step and fitted at the resolution of the two aneuploid subpopulations according to 6.2.
